# Differential mosquito attraction to humans is associated with skin-derived carboxylic acid levels

**DOI:** 10.1101/2022.01.05.475088

**Authors:** Maria Elena De Obaldia, Takeshi Morita, Laura C. Dedmon, Daniel J. Boehmler, Caroline S. Jiang, Emely V. Zeledon, Justin R. Cross, Leslie B. Vosshall

## Abstract

Female *Aedes aegypti* mosquitoes feed on human blood, which they use to develop their eggs. It has been widely noted that some people are more attractive to mosquitoes than others, but the mechanistic basis of this phenomenon is poorly understood. Here we tested mosquito attraction to skin odor collected from human subjects and identified people who are exceptionally attractive or unattractive to mosquitoes. Notably, these preferences were stable over several years, indicating consistent longitudinal differences in skin odor between subjects. We carried out gas chromatography/quadrupole time of flight-mass spectrometry to analyze the chemical composition of human skin odor in these subjects and discovered that highly attractive people produce significantly increased levels of carboxylic acids. Mosquitoes could reliably distinguish a highly attractive human from their weakly attractive counterparts unless we substantially diluted the odor of the “mosquito magnet.” This is consistent with the hypothesis that odor concentration is a major driver of differential attraction, rather than the less-favored skin odor blend containing repellent odors, although these are not mutually-exclusive. Mosquitoes detect carboxylic acids with a large family of odor-gated ion channels encoded by the Ionotropic Receptor gene superfamily. Mutant mosquitoes lacking any of the Ionotropic Receptor (IR) co-receptors *Ir8a, Ir25a,* and *Ir76b,* were severely impaired in attraction to human scent but retained the ability to differentiate highly and weakly attractive people. The link between elevated carboxylic acids in “mosquito-magnet” human skin odor and phenotypes of genetic mutations in carboxylic acid receptors suggests that such compounds contribute to differential mosquito attraction. Understanding why some humans are more attractive than others provides insights into what skin odorants are most important to the mosquito and could inform the development of more effective repellents.

## INTRODUCTION

The globally invasive mosquito *Aedes aegypti* is a highly efficient vector of viruses including yellow fever, dengue, chikungunya, and Zika among human populations (*1*). A single female mosquito will bite multiple humans during her 3 to 6-week lifetime to obtain sufficient protein to produce a new batch of eggs as often as every four days (*2*). This repetitive human-directed feeding behavior allows the mosquito to contract and transmit pathogens in successive bites. An important factor in the effectiveness of *Aedes aegypti* as a disease vector is that it specializes on human hosts (*3–5*), thereby focusing pathogen transmission among our species. Female *Aedes aegypti* mosquitoes have a strong innate drive to hunt humans, using sensory cues including exhaled CO_2_, body heat, and skin odor. While CO_2_ and heat are generic stimuli that signify a living warm-blooded animal, skin odor provides information about whether the target is a human or non-human animal (*3–5*). It is well-documented that mosquitoes are more strongly attracted to some humans than others (*6–20*), but the underlying mechanisms for this phenomenon remain unclear. The observation that some people are “mosquito magnets” is a topic that captivates the general public and scientific community alike. There is much speculation about possible mechanisms, but only some have a scientific basis. A common explanation offered by non-experts is that differences in ABO blood type “explain” attractiveness to mosquitoes, but experimental data that address this belief are contradictory (*21–25*). The widely quoted efficacy of eating garlic (*26*) or B vitamins (*27*) as a home remedy to repel mosquitoes is similarly unclear. Although a twin study documented a strong heritable component (*18*), non-genetic factors also contribute to selective attractiveness to mosquitoes. A given person can become more attractive to mosquitoes in contexts including pregnancy (*12, 28*), malaria parasite infection(*29–33*), and beer consumption (*34, 35*). The most widely accepted explanation for these differences is that variation in skin odors produced by different humans, related in part to their unique skin microbiota (*9, 17*), governs their attractiveness to mosquitoes (*8, 13–16*). However, the specific chemical mechanism for differential attractiveness to mosquitoes remains unclear.

Human skin odor is a blend of many organic compounds (*36–38*), the composition of which has not been exhaustively inventoried. It remains unclear how consistent human skin odor is over time within an individual. Whereas much work has focused on characterizing human axillary (armpit) malodor, there is relatively little information about the composition of the markedly less intense skin odor that emanates from body sites commonly bitten by mosquitoes. Furthermore, additional work is needed before we can fully appreciate the extent of interindividual variation in human skin odor. Thus, it is not known which specific components are most relevant for mosquito attraction to humans, nor do we understand which odorants cause mosquitoes to choose to bite some people over others. Humans who are highly attractive to mosquitoes may produce more attractant odors than other people (*5*). Alternatively, less attractive humans may emit compounds that repel mosquitoes (*39, 40*). To date no single molecule obtained from human skin can be said to be sufficient to explain how attractive a person is to mosquitoes. Blends of odorants can be more or less attractive depending on the composition of the blend and the concentration of a specific molecule. For example, the binary blend of ammonia and lactic acid strongly synergizes to elicit mosquito attraction (*41, 42*). Although carboxylic acids are neutral or repellent when presented individually or in combination with each other, they strongly increase mosquito attraction when combined with ammonia and lactic acid (*43, 44*). Mosquito attraction behavior is elicited much more reliably using live human hosts or natural odor blends collected directly from humans, than it is by mixing pre-specified compounds, despite improvements in synthetic odor blends as lures for attract-and-kill traps for use in the field (*45–47*). Moreover, the current absence of a complete reference metabolome of chemical compounds found on human skin, and the lack of commercially available standard molecules for many skin compounds also limits the effectiveness of human odor blend reconstitution approaches for studying mosquito attraction.

Mosquitoes use two large multigene families to detect olfactory cues that each encode odor-gated ion channels, the odorant receptors (ORs) and the ionotropic receptors (IRs) (*48–54*). ORs and IRs are evolutionarily unrelated, but both assemble into multi-subunit complexes with a ligand-selective subunit and a co-receptor subunit that does not respond to odorants (*48–54*). There are hundreds of ligand-selective ORs and IRs in a given insect species, but only one OR co-receptor (*Orco*) and three IR co-receptors (*Ir8a*, *Ir76b*, *Ir25a*). In *Aedes aegypti* there are 116 ligand-selective ORs and 132 ligand-selective IRs (*55*). Together these large gene families of odor-gated ion channels sense a vast number of chemical ligands. Although there is some overlap in ligand tuning, ORs generally respond to esters, alcohols, ketones, and aldehydes, and IRs respond to carboxylic acids and amines (*50, 56*). Because of this co-receptor organization, mutating a single co-receptor gene leads to profound deficits in the ability of an insect to detect whole classes of odorants (*49, 54, 57, 58*). Nevertheless, mosquitoes are remarkably resilient in the face of such genetic manipulations. Animals lacking the major receptor for carbon dioxide, *Gr3*, continue to be attracted to humans in semi-field conditions (*59*). Mosquitoes with a loss of function mutation in *Orco* lose strong preference for humans over non-human animals but retain strong attraction to humans overall (*60, 61*). Finally, *Ir8a* mutants show severe deficits in detecting lactic acid, a major human skin odor, but nevertheless remain partially attracted to humans (*61*). The recent discovery of extensive co-expression of ORs and IRs in single olfactory sensory neurons may explain this functional redundancy (*62, 63*).

In this paper we analyzed the skin-derived compounds that differentiate highly from weakly attractive humans and asked which mosquito sensory pathways are required to distinguish such people. We developed a new two-choice behavioral assay that allowed us to test mosquito attraction with higher throughput, allowing for frequent, repeat sampling of human subjects. We collected human skin odor samples on nylon stockings worn on the forearms and profiled the attractiveness of 64 human subjects to mosquitoes. We identified a cohort of highly and weakly attractive people and discovered that the *Orco* co-receptor is not required for discriminating between them. Mutants lacking *Ir8a*, *Ir76b*, and *Ir25a* retained a preference for “mosquito magnets,” but showed an overall reduced attraction to human skin odor (*61*). Therefore although neither the OR nor the IR pathway is solely required for discriminating among different people, mutating the IR pathway produced significantly stronger effects on overall mosquito attraction to humans than the OR pathway. We used gas chromatography/quadrupole time of flight-mass spectrometry (GC/QTOF-MS) to identify skin odor molecules that are associated with attractiveness to mosquitoes. Because carboxylic acids have been shown to be attractive to mosquitoes (*44*), we focused our chemical analysis on detecting acids in the human skin odor blend by using specialized sample preparation to enrich for highly polar acids. Since these are otherwise difficult to detect using standard analytical chemistry approaches, their contribution to differential mosquito attraction to humans is relatively understudied (*38*). We determined that highly attractive humans have higher levels of several carboxylic acids on their skin than less attractive humans. When we substantially diluted nylons from the most highly attractive subject, mosquitoes were no longer able to distinguish this subject from the least attractive subjects. Our results strongly suggest a link between the known function of IRs in acid-sensing and our observation that skin-derived carboxylic acids are associated with a person being a “mosquito magnet.”

## RESULTS

### Mosquitoes show strong preferences for individual humans

Preferential mosquito attraction to different stimuli is typically measured in a two-choice olfactometer. In previous studies, we used a Gouck olfactometer (*64*) to characterize female *Aedes aegypti* preferences for a human or non-human animal (*5, 60*). This large apparatus was not suitable for the higher-throughput analysis of mosquito preference for humans required for this study. We therefore adapted a previously described single-stimulus olfactometer (*65*) to reconfigure it as a two-choice olfactometer (Figure 1A), allowing us to test mosquito preferences between the forearms of two different live human subjects or their forearm skin odor collected on nylon sleeves. In this assay, a mixture of air and carbon dioxide (CO_2_) was passed over each stimulus to convey volatile odors to mosquitoes downwind (Figure 1A). Mosquitoes flew upwind and those that entered either of the two cylindrical traps in front of each stimulus were scored as attracted. The two-choice assay tests mosquito attraction to odor stimuli that are ∼0.91 m away, meaning that to reach the attraction trap, mosquitoes must travel ∼610 times their body length, assuming an average female mosquito thorax length of 1.5 mm (*66*). During the 3.1-year course of this study, we carried out more than 2,330 behavior trials on 174 experimental days. In control experiments we verified that there was no difference in attraction when mosquitoes were offered the left and right forearms of the same subject (Figure 1B). In pilot experiments comparing mosquito attraction among all possible pairings of 3 live human subjects, we identified one (Subject 33) that was significantly more attractive than two others (Subjects 25 and 28) (Figure 1B). Moreover, when the two less attractive subjects were competed against each other, Subject 25 attracted significantly more mosquitoes than Subject 28 (Figure 1B).

**Figure 1:**
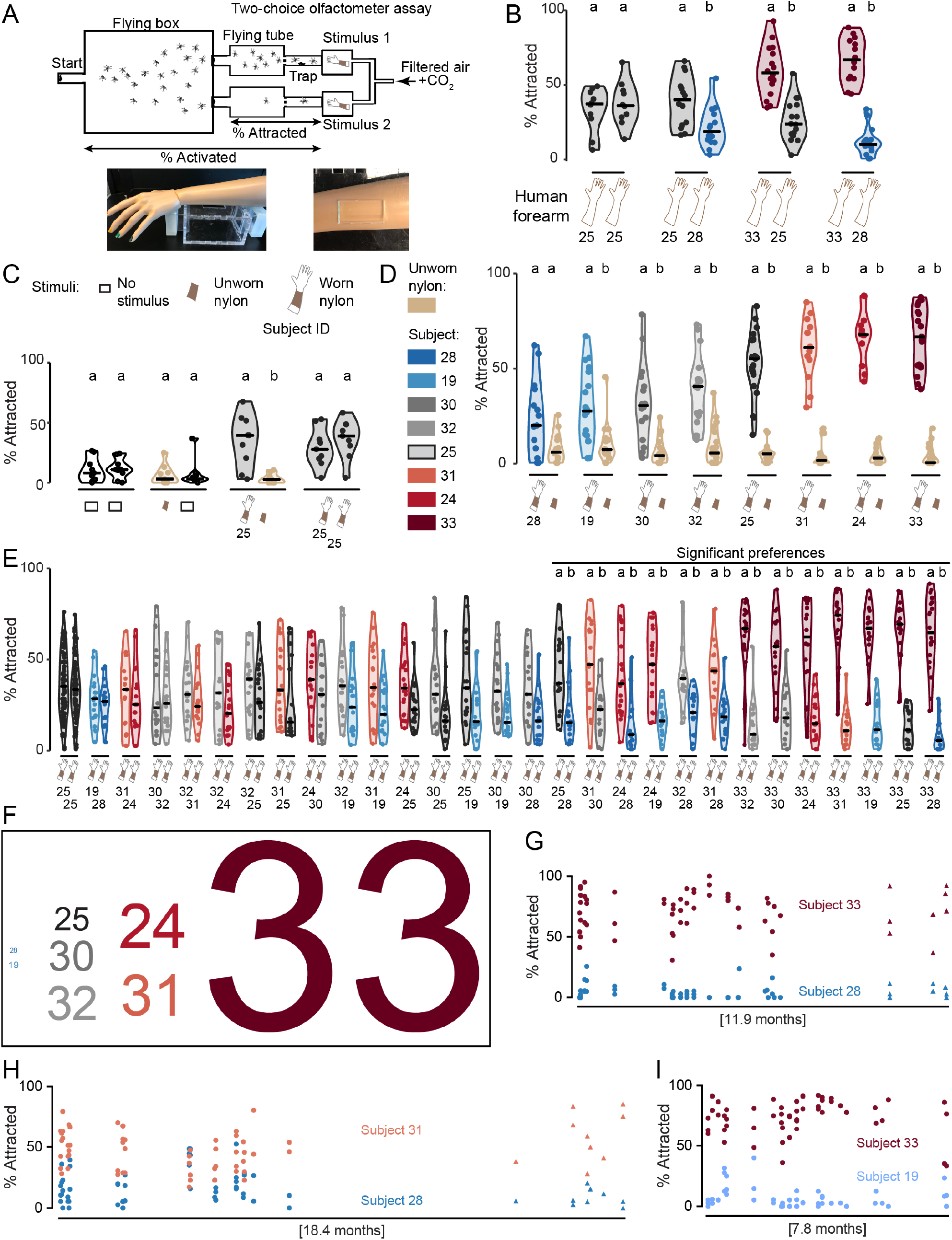
Mosquitoes show strong preferences among individual humans. (**A**) Schematic of two-choice olfactometer assay (top). Photographs (bottom) of a mannequin arm modeling the position of a live human forearm on top of the stimulus box in the two-choice olfactometer (left) and the opening in the stimulus box lid used to expose an area of human skin (5.08 cm x 2.54 cm) in the assay (right). (**B-E**) Wild-type mosquitoes attracted to live human forearms (B) or a 5.08 cm x 2.54 cm piece of human-worn nylons and controls (C-E) of the indicated subjects in the two-choice olfactometer assay. Subject pairs in E are ordered by nonparametric effect size. **(F)** Attractiveness scores for human subjects derived from data in E, with subject ID font size scaled to the attractiveness score: Subject 33 (score=144), Subject 24 (score=34), Subject 31 (score=32), Subject 32 (score=26), Subject 30 (score=23), Subject 25 (score=18), Subject 19 (score=1), Subject 28 (score=0). **(G-I)** Longitudinal two-choice olfactometer data for two wild-type *Aedes aegypti* mosquito strains, Orlando (circles) and Liverpool (triangles), showing attraction to the indicated subject pairs. The total time elapsed between the first and last experiment shown is indicated, and corresponded to July 12, 2018 to July 3, 2019 (G), February 1, 2018 to August 6, 2019 (H), and July 30, 2018 to March 21, 2019 (I). In B-E, data are displayed as violin plots with median indicated by horizontal black lines and the bounds of the violin corresponding to the range (30-40 mosquitoes/trial). B: n=10-16 trials, C: n=7-8 trials, D: n=12-21 trials, E: n=14-20 trials (except n=81 trials for the Subject 25 vs 25 comparison). Data corresponding to adjacent violin plots labeled with different letters are significantly different (p<0.05, Wilcoxon rank-sum tests with Bonferroni correction).

The two-choice olfactometer as configured for live human subjects requires participants to be physically present for competitions that take place in a warm, humid room. To make it feasible to carry out hundreds of competitions between two subjects over many months, we turned to using human forearm odor collected on nylon sleeves as previously described (*5, 60*). Empty stimulus traps did not attract mosquitoes, and mosquitoes did not prefer unworn nylon over an empty trap (Figure 1C). However, a 3” x 4” swatch cut from nylons worn by Subject 25 was significantly more attractive than the same-sized swatch from an unworn nylon (Figure 1C). To check for side bias in the assay, we used nylons from Subject 25 in both stimulus traps and confirmed that mosquitoes showed no preference (Figure 1C).

To study interindividual differences in attractiveness to mosquitoes, we recruited an additional 5 human subjects who provided skin-scented nylon samples frequently over a period of several months. When we tested nylons worn by each of these 8 subjects against unworn nylons in the two-choice olfactometer assay, we found that each was significantly more attractive than an unworn nylon (Figure 1D). However, there were remarkable differences in the attractiveness of the 8 subjects. When we ranked median attraction, we found that nylons from Subject 28 attracted few mosquitoes, Subject 25 showed intermediate attractiveness, whereas Subject 33 was highly attractive (Figure 1D), mirroring the results of the live human experiments in Figure 1B. This suggests that skin odor is the primary driver of differential mosquito attraction to humans, since temperature and CO_2_ cues were held constant across all of our nylon experiments. It also suggests that skin odor captured on forearm-worn nylon sleeves is a good approximation of the odor emanating from a live human forearm. When we tested the 5 additional subjects, we found one additional low attractor (Subject 19) and two additional high attractors (Subjects 24 and 31). Two other subjects showed intermediate attractiveness (Subjects 30 and 32) (Figure 1D).

We reasoned that in a real-world situation, mosquitoes would choose among multiple different humans in a local area, such that the absolute attractiveness of a single human would not necessarily predict their attractiveness relative to another person. To systematically determine the relative attractiveness of these 8 humans to mosquitoes, we performed a round-robin style “tournament”, competing nylons from all possible subject pairings from this group of 8 subjects, for a total of 28 separate competitions using the two-choice olfactometer assay (Figure 1E). On each experimental day, we verified that mosquitoes did not show a preference when both stimulus ports were baited with nylons collected from the same subject. We sampled each pair of humans on 6 separate days over a period of several months (558 trials, performed over 42 experimental days). Among 28 subject pairs tested, we found 13 pairs for which mosquitoes significantly preferred one subject’s odor over the other (Figure 1E). In the other cases, mosquitoes had no preference for one subject over the other. Subject 33 attracted significantly more mosquitoes than every other subject in essentially every trial performed, usually by a large margin. Subjects 19 and 28 were significantly less attractive than several other subjects. When mosquitoes were presented with a choice between the two low attractors, Subjects 19 and 28, they were not able to distinguish between them (Figure 1E). To rank subjects from most to least attractive, we devised a scoring system (“attraction score”) based on how many more mosquitoes each subject attracted when competed against all 7 other subjects. By this metric, Subject 33 was the most attractive, yielding an attractiveness score that was 4 times the attractiveness score of the next most attractive subject, and over 100 times greater than that of the two least attractive Subjects 19 and 28 (Figure 1F). These differences in attraction to specific pairs of humans were remarkably stable over many months and were seen with two different wild-type strains of *Aedes aegypti* (Figure 1G-I). Taken together these results provide empirical evidence that mosquitoes strongly prefer some people over others, and that the olfactory cues that make some people mosquito magnets are stable over many months.

### *Orco* and *Ir8a* mutant mosquitoes retain individual human preferences

We have shown that small swatches of human-scented nylon provide enough information for mosquitoes to distinguish between and prefer one person over another. What sensory mechanisms do mosquitoes rely on to detect these interindividual differences in skin odor? We first tested the preference of mosquitoes lacking the OR co-receptor *Orco*, which retain strong attraction to humans, but show severe deficits in discriminating humans from non-human animals (*60*). *Orco* mutant mosquitoes showed wild-type levels of activation and attraction in control trials where nylons from Subject 25 were placed in both stimulus boxes (Figure 2A). In the course of carrying out these control experiments we noted low participation in some trials. Therefore, we put inclusion criteria in place such that trials in which 9 or fewer mosquitoes entered either trap were excluded. We note that low levels of participation preclude the accurate calculation of attraction preferences, such as an instance in which only 3 mosquitoes entered either trap. A high percentage of excluded trials may reflect an overall deficit in locomotor activity or general decreases in activation and attraction to human sensory cues or a combination of both factors.

**Figure 2:**
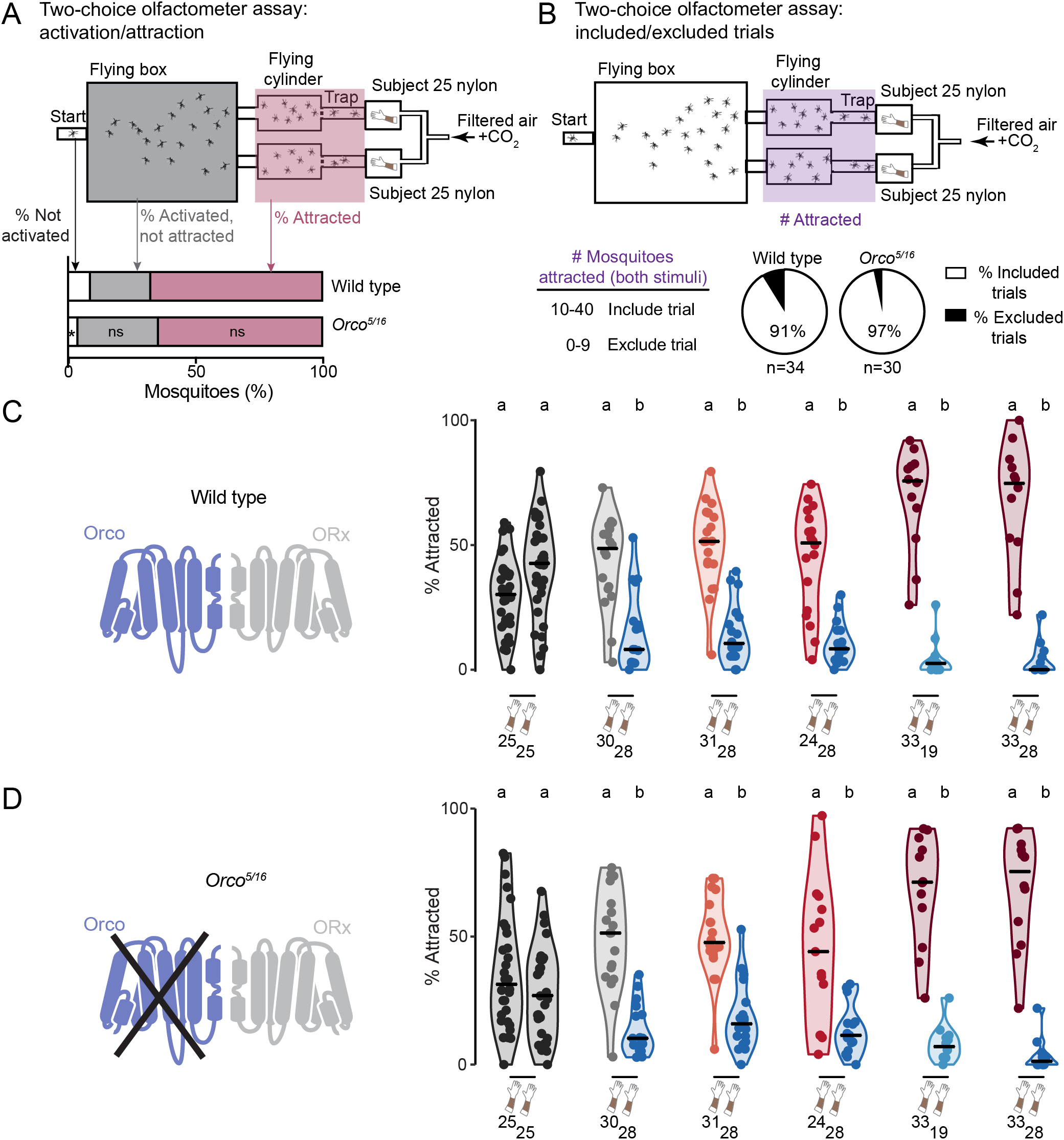
Mosquitoes lacking *Orco* retain individual human preferences. (A) Schematic of two-choice olfactometer assay indicating the location of mosquitoes that were not activated, activated but not attracted, or attracted in response to a control trial in which Subject 25 nylons were placed in both stimulus boxes. Stacked bar plots indicate the mean total percent of mosquitoes that were in each category in all trials (30-40 mosquitoes/trial, n=30-34 trials, *p<0.01, Wilcoxon rank-sum tests with Bonferroni correction comparing each category across the two genotypes). (B) Top: schematic of two-choice olfactometer assay, indicating the location (purple shading) of all mosquitoes attracted to either stimulus in a control trial in which Subject 25 nylons were placed in both stimulus boxes. Bottom: trials in which 9 or fewer animals entered either trap were excluded. (**C,D**) Left: schematic of Orco and ligand-specific subunit (ORx). Right: percent of mosquitoes of the indicated genotype attracted to the indicated stimuli in the two-choice olfactometer assay. Data from trials that met the inclusion criteria are displayed as violin plots with median indicated by horizontal black lines and the bounds of the violin corresponding to the range (30-40 mosquitoes/trial, n=11-18 trials, except n=29-31 for the Subject 25 vs 25 comparison). Data corresponding to adjacent violin plots labeled with different letters are significantly different (p<0.05, Wilcoxon rank-sum tests with Bonferroni correction).

More than 90% of all such control trials using both wild type and *Orco* mutants resulted in at least 10 total mosquitoes being attracted to either stimulus (Figure 2B). Among trials that met the inclusion criteria, *Orco* mutants did not differ from wild-type controls, retaining the ability to distinguish 5 pairs of highly and weakly attractive humans (Figure 2C-D). This is an important result because in *Orco* mutants none of the 117 OR genes (*55*) are functional, suggesting that either ligands that activate ORs do not contribute to interindividual differences in human attractiveness to mosquitoes or that mosquitoes have redundant chemosensory abilities to detect these differences.

We next tested mosquitoes lacking the co-receptor *Ir8a*, which is expressed in the antenna and necessary for detection of several acids, including lactic acid, a component of human sweat (*61*). *Ir8a* mutants showed decreased overall attraction to Subject 25 in the two-choice olfactometer assay, despite normal levels of activation, as defined by entry into the flying chamber (Figure 3A). In control trials using Subject 25 nylons in both stimulus boxes, 94% of trials using wild-type mosquitoes resulted in 10 or more total mosquitoes being attracted to either stimulus, but this was reduced to 73% for trials using *Ir8a* mutants (Figure 3B). Among trials that met the inclusion criteria, *Ir8a* mutants largely retained the same preferences as wild-type controls, despite their diminished overall attraction to human odor (Figure 3C-D). This result again suggests that mosquitoes impaired in detecting specific human body odors are still able to tell the difference between different people.

**Figure 3:**
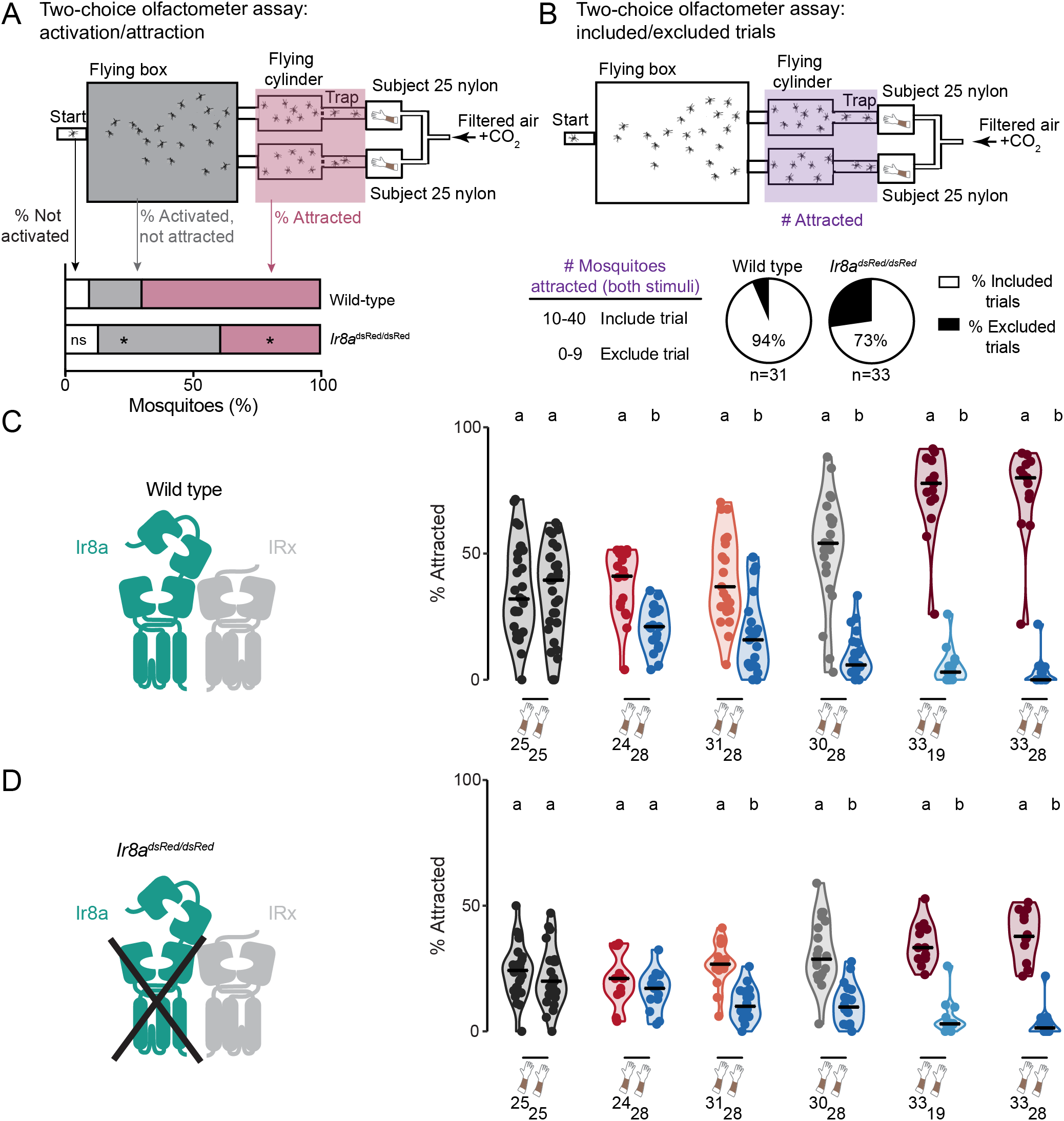
Mosquitoes lacking *Ir8a* retain individual human preferences. (A) Schematic of two-choice olfactometer assay indicating the location of mosquitoes that were not activated, activated but not attracted, or attracted in response to a control trial in which Subject 25 nylons were placed in both stimulus boxes. Stacked bar plots indicate the mean total percent of mosquitoes that were in each category in all trials (30-40 mosquitoes/trial, n=31-33 trials, *p<0.0001, Wilcoxon rank-sum tests with Bonferroni correction comparing each category across the two genotypes). (B) Top: schematic of two-choice olfactometer assay, indicating the location (purple shading) of all mosquitoes attracted to either stimulus in a control trial in which Subject 25 nylons were placed in both stimulus boxes. Bottom: trials in which 9 or fewer animals entered either trap were excluded. (**C, D**) Left: Schematic of Ir8a and ligand-specific subunit (IRx). Right: percent of mosquitoes of the indicated genotype attracted to the indicated stimuli in the two-choice olfactometer assay. Data from trials that met the inclusion criteria are displayed as violin plots with median indicated by horizontal black lines and the bounds of the violin corresponding to the range (30-40 mosquitoes/trial, n=11-22 trials, except n=24-29 for the Subject 25 vs 25 comparison). Data corresponding to adjacent violin plots labeled with different letters are significantly different (p<0.05, Wilcoxon rank-sum tests with Bonferroni correction).

### Generation and behavioral characterization of *Ir76b* and *Ir25a* mutants

We next used CRISPR-Cas9 to generate mosquitoes that lack the two other IR co-receptors *Ir76b* and *Ir25a* (Figure 4A-B, Supplementary Figure S1). We generated 2 mutant alleles of each gene (*Ir76b^32^*, *Ir76b^61^*, *Ir25a^BamHI^*, and *Ir25a^19^*) and tested the behavior of heterozygous animals of all 4 strains and the heteroallelic *Ir76b^32/61^* and *Ir25a^BamHI/19^* null mutants. We noted that homozygous *Ir76b* and *Ir25a* mutants had difficulty blood-feeding and that *Ir25a* homozygous mutants generally laid fewer eggs. A recent analysis of *Anopheles coluzzii Ir76b* null mutants reported that they show normal attraction to human host cues but do not blood-feed or produce eggs (*67*). To characterize the response of these strains and wild-type control mosquitoes to human cues, we first used a single-stimulus olfactometer (*65*) to examine activation and attraction levels (Figure 4C), before conducting preference assays. *Ir76b^32/61^* mutants displayed reduced general activity levels in response to an unworn nylon or a nylon worn by Subject 33 (Figure 4D). However, this defect was readily overcome when *Ir76b* mutants were presented with the forearm of Subject 33, indicating the absence of gross motor defects (Figure 4D). *Ir25a^BamHI/19^* mutants showed wild-type levels of activation across all stimuli tested (Figure 4E). We additionally asked if the *Ir76b* and *Ir25a* mutants were attracted to the same stimuli described above. *Ir76b^32/61^* mutants showed normal levels of attraction to nylons worn by Subject 33 and to the forearm of Subject 33 (Figure 4F). In contrast, *Ir25a^BamHI/19^* mutants displayed significant defects in their attraction to both Subject 33 nylons and to the forearm of Subject 33 (Figure 4G). This suggests that the *Ir25a* co-receptor, along with one or more of the ligand-selective IRs with which it assembles a functional receptor, plays an important role in detecting human skin emanations.

**Figure 4:**
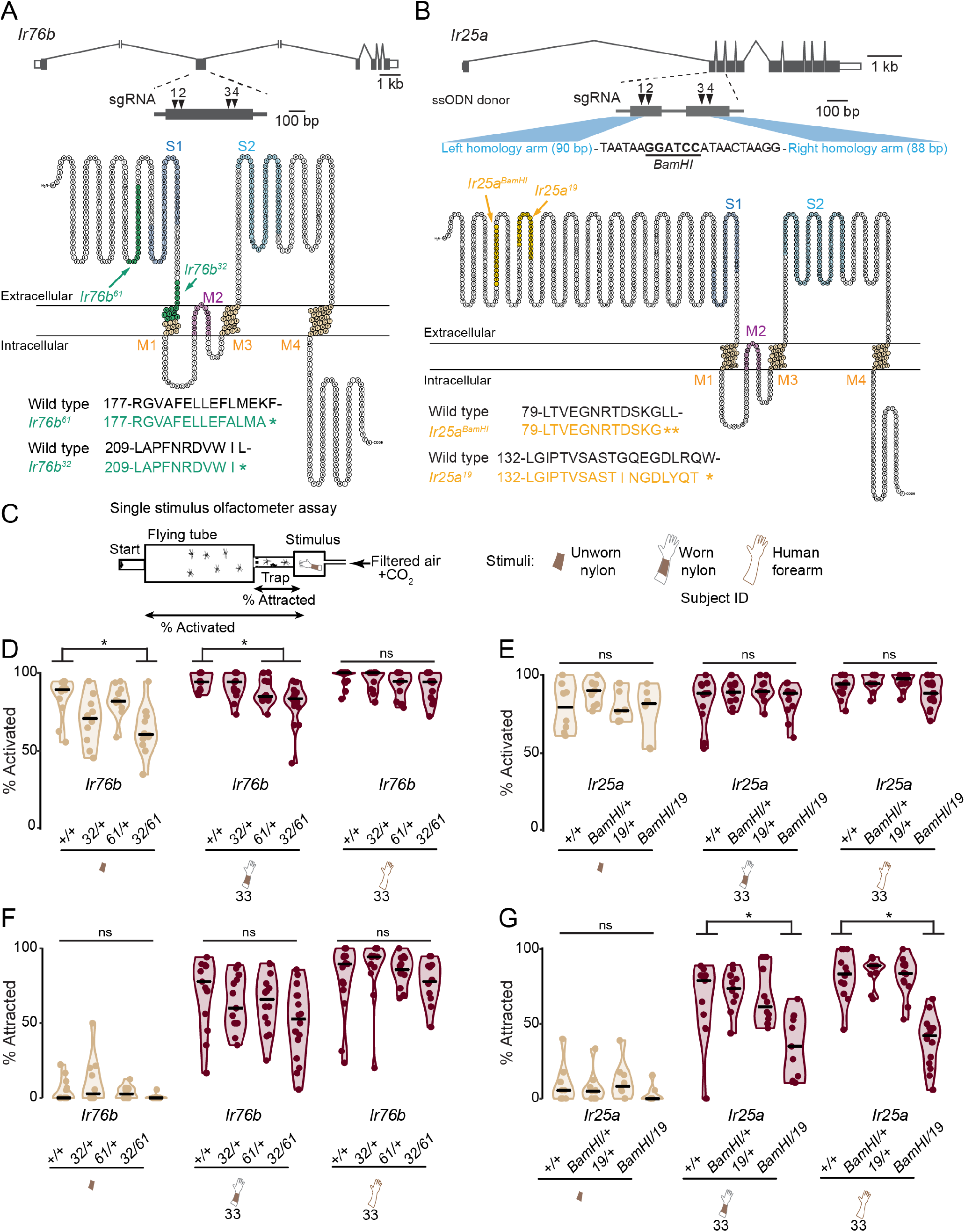
Generation and characterization of *Ir25a* and *Ir76b* mutants. (**A-B**) Schematic of the *Aedes aegypti Ir76b* (A) and *Ir25a* (B) genomic loci, detailing sgRNA sites and modified protein products of the indicated mutant alleles superimposed on Ir76b and Ir25a protein snake plots, which were generated using Protter v1.0 (*84*). (C) Schematic of single stimulus olfactometer assay. (**D-G**) Percent of mosquitoes of the indicated genotypes activated to leave the start canister (D-E) or attracted to the indicated stimuli (F-G) in the single stimulus olfactometer assay. Data are displayed as violin plots with median indicated by horizontal black lines and the bounds of the violin corresponding to the range (10-20 mosquitoes/trial, n=6-16 trials). Kruskal-Wallis test was used to compare each mutant allele to wild-type controls (ns, not significant; *p<0.05).

### *Ir76b* and *Ir25a* mutant mosquitoes retain individual human preferences

Given the dramatic decrease in attraction of *Ir25a^BamHI/19^* mutant mosquitoes to Subject 33, we next asked if these mutants along with *Ir76b^32/61^* mutants could distinguish between nylons worn by highly and weakly attractive human subjects, when these were presented simultaneously in the two-choice olfactometer assay. The two-choice assay is substantially larger than the single-choice assay, and thus may represent a more difficult behavioral task, so we expected that it might reveal additional phenotypes not seen in Figure 4D-G.

In control two-choice assay trials in which Subject 25 nylons were placed in both stimulus boxes, *Ir76b^32/61^* mutants showed lower activation in response to human worn nylons, and both *Ir76b^32/61^* and *Ir25a^BamHI/19^* mutants showed significantly decreased attraction compared to wild-type controls (Figure 5A). Furthermore, only 25% of *Ir76b^32/61^* and 38% of *Ir25a^BamHI/19^* mutant trials had at least 10 mosquitoes attracted to either stimulus, compared to 100% of wild type trials (Figure 5B).

**Figure 5:**
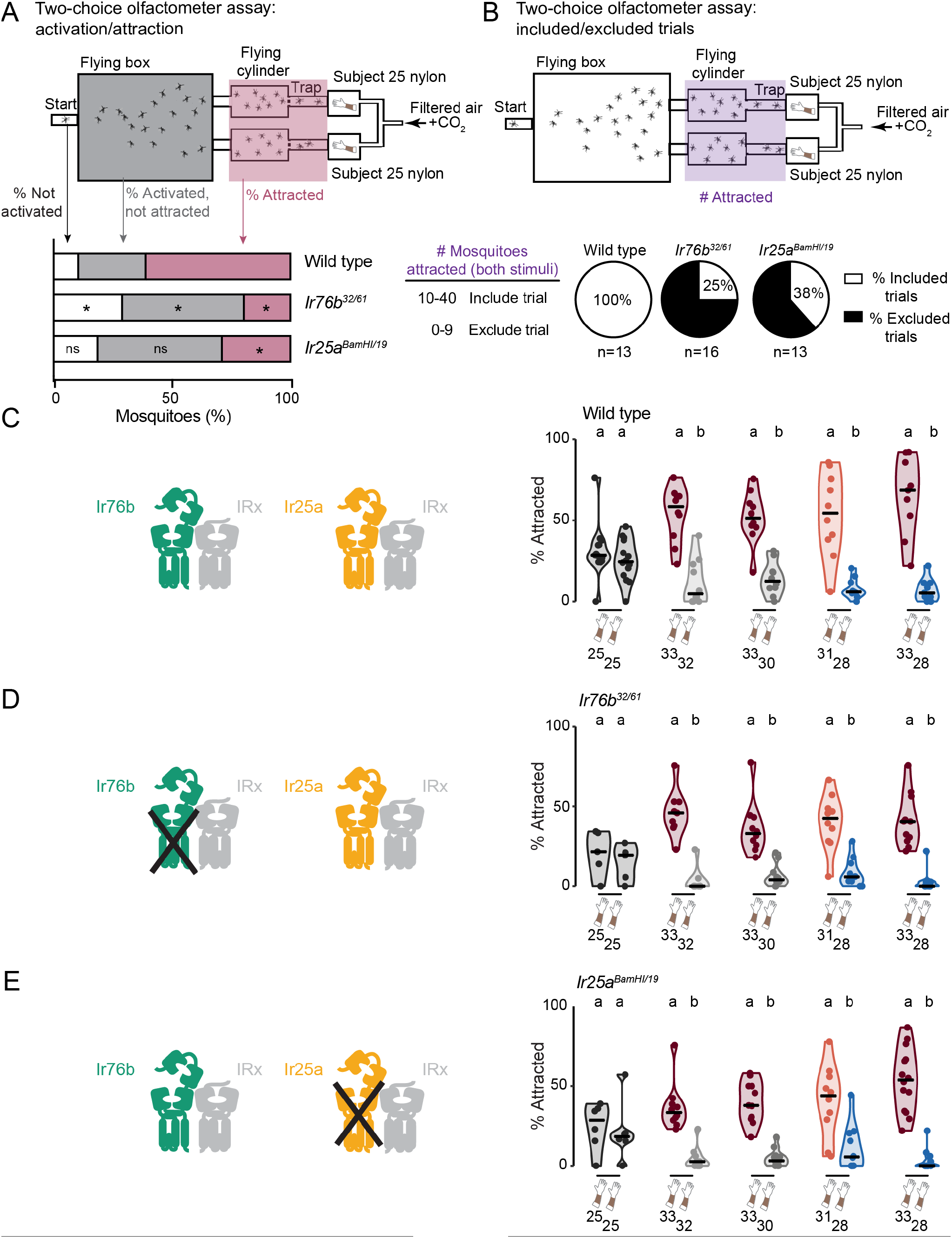
Mosquitoes lacking *Ir76b* or *Ir25a* show reduced attraction to humans but retain individual human preferences. (A) Schematic of two-choice olfactometer assay indicating the location of mosquitoes that were not activated, activated but not attracted, or attracted in response to a control trial in which Subject 25 nylons were placed in both stimulus boxes. Stacked bar plots indicate the mean total percent of mosquitoes that were in each category (30-40 mosquitoes/trial, n=13-16 trials, *p<0.05, Wilcoxon rank-sum tests with Bonferroni correction comparing each category across the two genotypes). (B) Top: schematic of two-choice olfactometer assay, indicating the location (purple shading) of all mosquitoes attracted to either stimulus in a control trial in which Subject 25 nylons were placed in both stimulus boxes. Bottom: trials in which 9 or fewer animals entered either trap were excluded. (**C-E**) Left: Schematic of Ir76b and Ir25a and ligand-specific subunit (IRx). Right: percent of mosquitoes of the indicated genotype attracted to the indicated stimuli in the two-choice olfactometer assay. Data from trials that met the inclusion criteria are displayed as violin plots with median indicated by horizontal black lines and the bounds of the violin corresponding to the range (30-40 mosquitoes/trial, n=8-13 trials, except n=4-13 for the Subject 25 vs 25 comparison). Data corresponding to adjacent violin plots labeled with different letters are significantly different (p<0.05, Wilcoxon rank-sum tests with Bonferroni correction).

Analysis of trials that met these inclusion criteria showed that despite substantial defects in overall attraction to human odor, both *Ir76b^32/61^* and *Ir25a^BamHI/19^* mutants were able to distinguish highly and weakly attractive human subjects (Figure 5C-E). These data indicate that mosquitoes have evolved highly redundant sensory systems permitting them to retain attraction to humans even with significant genetic disruption of their olfactory system (*62*). Nevertheless, mutating the IR pathway produced significantly stronger effects on overall mosquito attraction to humans than the OR pathway.

### Carboxylic acids are elevated in skin odor of highly attractive humans

We have shown that human-worn nylon sleeves provide the sensory cues needed for mosquitoes to reliably discriminate between humans. We reasoned that the chemistry of skin-derived odors in these nylons would provide insights into what distinguishes a highly attractive person from a weakly attractive person. We used gas chromatography/quadrupole time of flight-mass spectrometry (GC/QTOF-MS) to identify compounds on the nylon sleeves that were associated with mosquito attractiveness (Figure 6A). Because IR co-receptor mutants showed significantly reduced attraction to humans, we focused our chemical analysis on acidic compounds, which are detected by the IR pathway (*61, 68, 69*). We therefore did not investigate whether “mosquito-magnet” humans produce elevated levels of other compounds not captured by our analytical strategy. We collected nylons from 7 of 8 human subjects in our initial subject cohort on 4 days spaced at least 1 week apart, and then performed 4 independent analyses (Experiments 1.1-1.4; Figure 6B). Briefly, sections of human-exposed nylon sleeves were extracted in 80:20 methanol:water. The resulting extract was derivatized with pentafluorobenzyl bromide (PFB-Br), an aqueous-compatible derivatization reagent that reacts with carboxylic acids, improving their chromatography performance and ionization efficiency (Supplementary Figure S2A). Samples were then analyzed by GC-MS using negative methane chemical ionization on a Q-TOF instrument, allowing formula prediction from the mass of detected ions. This strategy has been widely used to measure volatile and semi-volatile carboxylic acids but to our knowledge it has not previously been applied to examine skin metabolites in the context of mosquito preferences.

**Figure 6:**
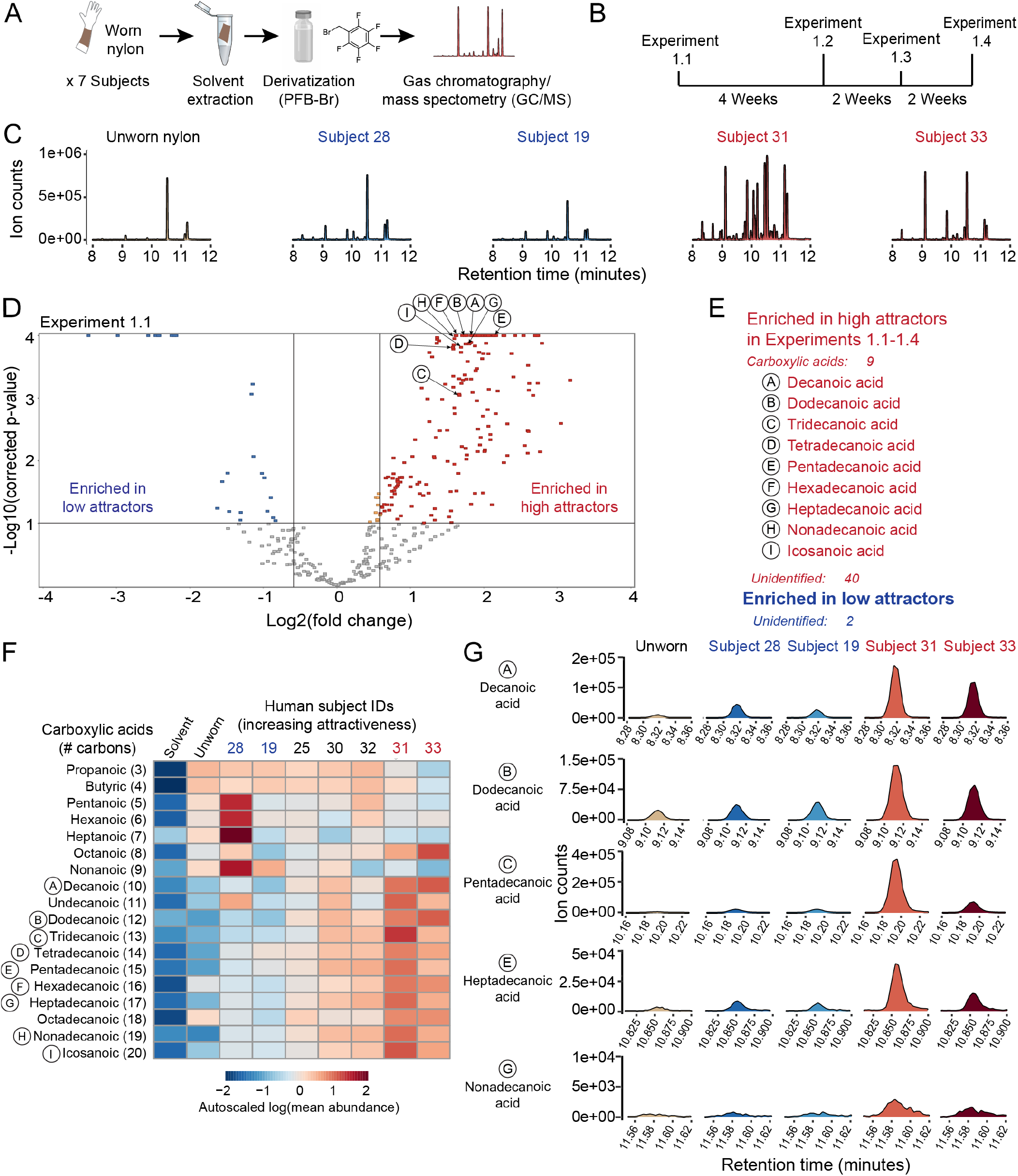
Carboxylic acids are enriched on the skin of humans who are highly attractive to mosquitoes. (A) Overview of experimental procedure for gas chromatography/quantitative time of flight mass spectrometry (GC/QTOF-MS) experiments. (B) Timeline of 4 replicate GC/QTOF-MS experiments in initial human subject cohort. (C) Representative chromatograms from the indicated sample groups, including merged extracted ion chromatograms from a set of ∼200 features enriched on worn nylons versus unworn nylons and solvent controls in Experiments 1.1-1.4 (Supplementary Table S1). (D) Volcano plot of features enriched on worn nylons versus unworn nylons and solvent controls in Experiment 1.1. Nine identified compounds that were differentially abundant between high and low attractor groups in Experiments 1.1-1.4 are indicated. (E) Table of differential features in Experiments 1.1-1.4. (F) Heatmap quantifying abundance of carboxylic acids with 3-20 carbons in the indicated human subjects, averaged across 4 experiments. (G) Representative extracted ion chromatograms of several carboxylic acids in the two most and least attractive subjects from the initial cohort.

Using unbiased feature detection and data filtering we identified 204 molecular features that were likely to be human-derived, since they were enriched on subject nylons versus unworn nylons and method blanks, across all 4 experiments (Supplementary Table S1). We noted that high attractor subjects appeared to have more of these putative “human-derived” peaks overall than low attractor subjects (Figure 6C). Of these, ∼50 features were differentially present in samples from the two most attractive subjects, Subjects 33 and 31, versus the two least attractive subjects, Subjects 19 and 28, in all 4 replicate experiments (Figure 6D,E, Supplementary Table S1). Interestingly, nearly all differentially enriched features (49/51) were more abundant in the 2 highly attractive subjects, although we did find 2 features that were enriched in the 2 low attractors (Figure 6E). We were able to predict chemical formulas for about 40 of these features and ultimately identified 9 as straight chain fatty acids by matching their mass and retention time to that of authentic standards (Figure 6E). We then extracted the signals for all the straight chain acids with acyl chain lengths between 3-20 carbons (Figure 6F). Consistent with the untargeted analysis described above, these compounds were all low or absent in unworn nylons and solvent controls, but only fatty acids with >10 carbons appeared to be enriched in the most highly attractive subjects (Figure 6F-G, Supplementary Figure S2B). To exclude these changes being attributable to variations in sample loading, derivatization or extraction efficiency we verified that several control compounds were similarly abundant in the high and the low attractor samples, including 2 deuterated internal standards present in the extraction solvent and an unknown “nylon-derived” entity that was present in all nylon containing-samples, and absent in solvent-alone controls (Supplementary Figure S2C,D).

### Association of elevated skin-derived carboxylic acids and mosquito attraction confirmed in a validation cohort

To confirm our finding that highly mosquito-attracting humans produced more abundant carboxylic acids on their skin, we enrolled 56 new human subjects in a validation study (Supplementary Figure S3A) and used the higher-throughput single-stimulus olfactometer assay to screen the attractiveness of nylons from a given new subject compared to an unworn nylon (Figure 7A). Alongside the 56 new subjects, we also tested nylons from 7 subjects from the initial cohort in this assay (Figure 7B). Consistent with our initial behavioral studies, performed over 1 year earlier, Subject 28 was the least attractive of all 64 human subjects that we tested over the course of the whole study, and Subject 33 was among the most attractive subjects (Figure 7B, Figure 1G). Based on initial behavioral results we moved 18 subjects (4 from the initial cohort and 14 from the validation cohort) forward to metabolite profiling with GC/QTOF-MS. These comprised 11 subjects that were highly attractive and 7 that were weakly attractive. Subjects provided 4 more odor samples, spaced 1 week apart, which were used for additional behavioral testing to confirm their high/low attractor status, and for GC/QTOF-MS analysis. In Figure 7B we plot all data collected from the low and high attractor groups that were included in the GC/QTOF-MS analysis. Behavioral data from 45 additional subjects, 42 subjects from the validation cohort and 3 subjects from the initial cohort, who were not included in the GC/QTOF-MS validation study because of sample size limitations are available on Zenodo (DOI: 10.5281/zenodo.5822539).

**Figure 7:**
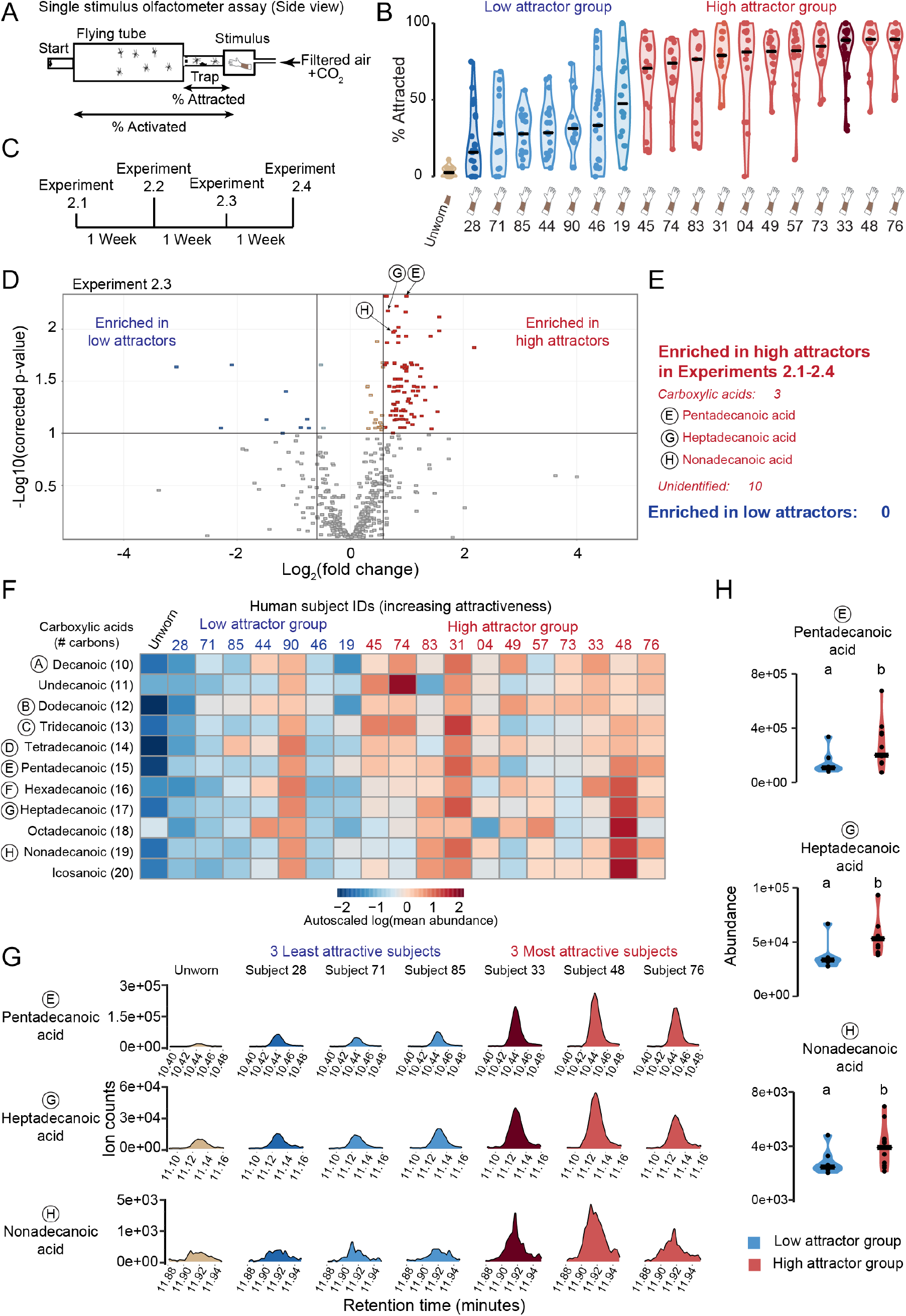
Carboxylic acids are enriched in a validation cohort of highly mosquito attractive humans. (A) Schematic of single stimulus olfactometer assay. (B) Mosquitoes attracted to nylons from 18 subjects at the extremes of low and high attraction in the single stimulus olfactometer assay, comprising 14 subjects from the GC/QTOF-MS validation study and 4 subjects from the initial cohort. Single stimulus olfactometer assay data from 45 additional subjects, comprising 42 subjects from the GC/QTOF-MS validation study and 3 subjects from the initial cohort are available on Zenodo (DOI: 10.5281/zenodo.5822539). Data are displayed as violin plots with median indicated by horizontal black lines and the bounds of the violin corresponding to the range (14-24 mosquitoes/trial, n=13-28 trials) (C) Timeline of 4 replicate GC/QTOF-MS experiments (Experiments 2.1-2.4), performed approximately one year after Experiments 1.1-1.4. (D) Volcano plot of features enriched on worn nylons versus unworn nylons and solvent controls in Experiment 2.3. Identified compounds that were differentially abundant between high and low attractors in all Experiments 2.1-2.4 are indicated with an arrow, and labeled with an uppercase letter, corresponding to the table in E. (E) Table describing features that were consistently differentially abundant in high versus low attractors in Experiments 2.1-2.4. (F) Heatmap quantifying abundance of carboxylic acids with 10-20 carbons, averaged across 4 experiments, in the 18 subjects in B. (G) Representative extracted ion chromatograms of three carboxylic acids in the 3 most and 3 least attractive subjects of the 18 subject validation cohort in B. (H) Quantified abundance (median peak areas) of three carboxylic acids in high attractors (n=11) versus low attractors (n=7) across Experiments 2.1-2.4. Data are displayed as violin plots with median indicated by horizontal black lines and the bounds of the violin corresponding to the range. Each plotted point represents the overall median abundance of the compound in one subject across Experiments 2.1-2.4. Data corresponding to adjacent violin plots labeled with different letters are significantly different (Wilcoxon rank-sum test followed by FDR correction p≤0.1).

We again performed 4 replicate metabolomic experiments (Figure 7C; Experiments 2.1-2.4) using PFB-Br derivatization to detect carboxylic acid-containing molecules. Since the validation study included more subjects than our initial cohort, we reasoned that untargeted analysis may reveal additional features of interest that were not found in the first study. Hence, we repeated the untargeted analysis of GC/QTOF-MS data, as described above, and found 161 molecular features that were likely to be human-derived, since they were more abundant in worn nylons from at least 1 subject than unworn nylons and solvent controls in all 4 experiments (Supplementary Table S1). We then filtered features that were differentially abundant in high versus low attractor groups in all 4 replicate experiments, resulting in a list of 13 features, all of which were enriched in the high attractor group (Figure 7D-E). We identified 3 of these features as carboxylic acids: pentadecanoic acid, heptadecanoic acid, and nonadecanoic acid (Figure 7E, Supplementary Figure S2G). We were not able to identify the remaining 10 features definitively, although in some cases we were able to predict a chemical formula. Of note, many of these features had the same predicted formula as the identified straight-chain fatty acids but eluted at different retention times, making it likely these are branched chain isoforms of the identified carboxylic acids. As expected, deuterated internal standards and a nylon-derived entity showed no difference in abundance between the high and low attractor groups (Supplementary Figure S2E,F).

We next performed a targeted re-analysis of carboxylic acids with 10-20 carbons in individual human subjects and control samples (Figure 7F-H). The abundance of carboxylic acids on individual subjects from the initial cohort was remarkably consistent with results obtained about 1 year earlier: low attractors Subjects 19 and 28 had much lower levels of many carboxylic acids than the more highly attractive Subjects 31 and 33 (Figure 6F, Figure 7F, Supplementary Figure S3B). The carboxylic acid pattern was similarly consistent from week to week for individual subjects in the larger cohort in Experiments 2.1-2.4 (Supplementary Figure S3C). Overall, the high attractor group had significantly higher levels of 3 carboxylic acids (pentadecanoic, heptadecanoic, nonadecanoic) than the low attractor subjects in this targeted re-analysis of the data (Figure 7G-H). However, not all individual subjects fit this pattern. Low attractor Subject 90 had high levels of all carboxylic acids examined, in contrast to the 6 other low attractors (Figure 7F). In both the initial and validation cohorts, we documented an association between high levels of skin carboxylic acids and attractiveness to mosquitoes.

### Dilution of highly attractive human odor eliminates mosquito preferences

Our GC/QTOF-MS experiments revealed that 3 carboxylic acids and 10 unknown compounds were consistently enriched in highly attractive subjects versus less attractive subjects, whereas we did not find any compounds that were consistently enriched in the low attractor group. Accordingly, we hypothesized that the high attractors have higher levels of mosquito attractant compounds on their skin. To test this hypothesis, we performed a dose-response experiment, in which we competed different sized swatches of high attractor Subject 33 nylon against a standard 5.08 cm x 2.54 cm-sized swatch of low attractor worn nylon from Subject 19 or Subject 28 (Figure 8). We found that mosquitoes preferred the odor blend of high attractor Subject 33 to that of both low attractor subjects, even when presented with a substantially smaller swatch of Subject 33 nylon than the less attractive subject’s nylon (Figure 8). Subject 33 nylon only became indistinguishable from low attractor nylons from Subject 28 and Subject 19 nylon, when competed against a swatch of Subject 33 nylon that was 32-fold or 8-fold smaller, respectively (Figure 8). Therefore, the skin odor blend of highly attractive Subject 33 provided a much more potent attractive stimulus than that of weakly attractive humans, which is consistent with our GC/QTOF-MS findings that highly attractive subject nylons contained more human-derived compounds, including carboxylic acids, than those of less attractive subjects.

**Figure 8:**
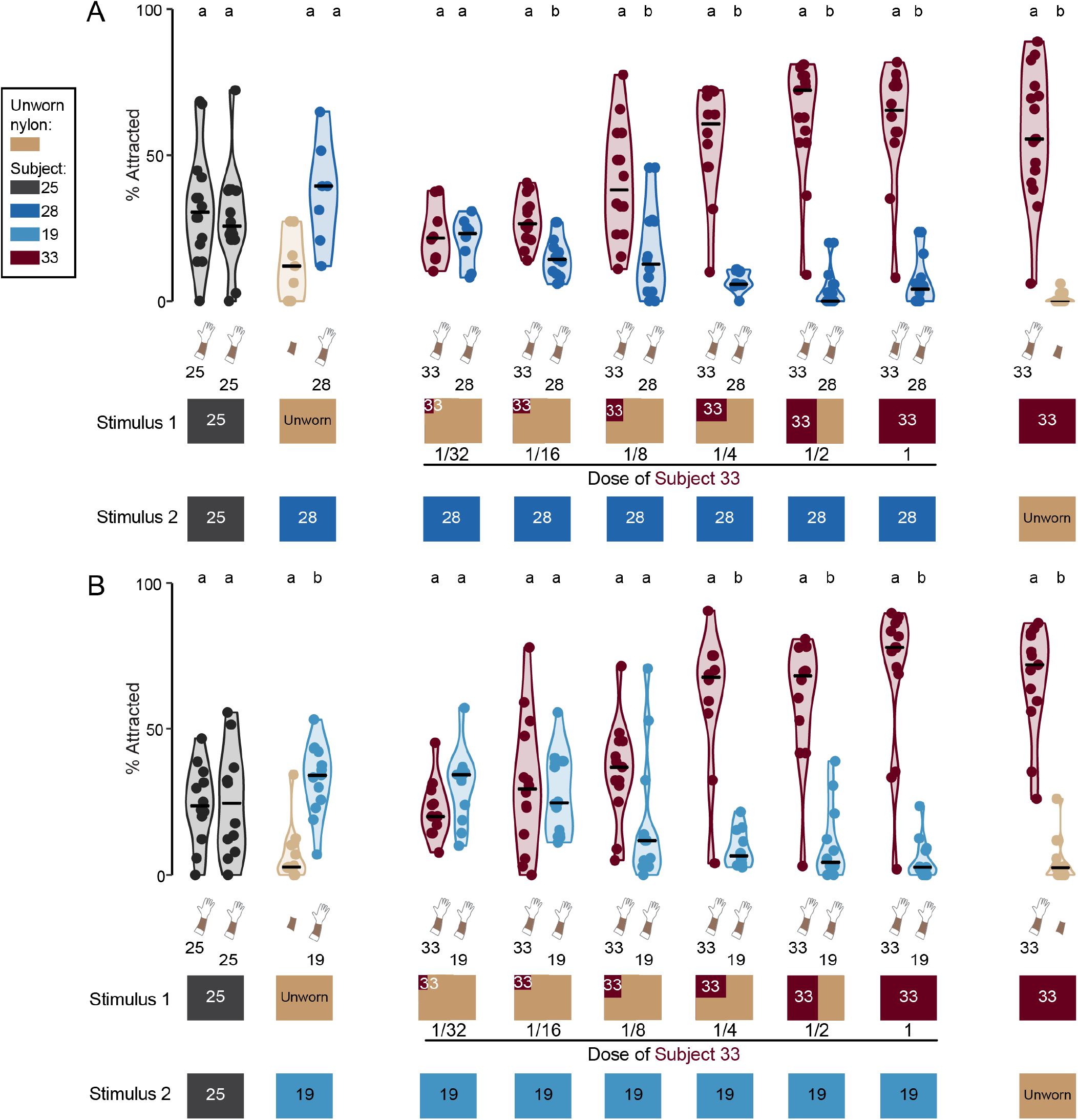
Dilution of highly attractive human odor eliminates mosquito preferences. (**A,B**) Percent of mosquitoes attracted to the indicated stimuli in the two-choice olfactometer assay. Mosquitoes were presented with a constant size of nylon worn by a low attractor, either Subject 28 (A) or Subject 19 (B), and decreasing amounts of nylon worn by high attractor Subject 33, corresponding to the indicated fraction of the low attractor nylon size. The total amount of nylon was balanced by adding unworn nylon. Data are displayed as violin plots with median indicated by horizontal black lines and the bounds of the violin corresponding to the range (30-40 mosquitoes/trial, n=11-20 trials). Data corresponding to adjacent violin plots labeled with different letters are significantly different (p<0.05, Wilcoxon rank-sum tests with Bonferroni correction).

## DISCUSSION

### Why are some people more attractive to mosquitoes than others?

In this work, we establish that the differential attractiveness of individual humans to mosquitoes is a stable over many months and is associated with the abundance of skin-associated carboxylic acids. Previous work demonstrated that mice at a specific stage of infection with malaria parasites were more attractive to mosquitoes, and that these infected mice showed an overall increase in concentration of many emitted volatile compounds (*32*). These results are consistent with our work. Highly attractive subjects produced significantly higher levels of 3 carboxylic acids—pentadecanoic, heptadecanoic, and nonadecanoic acids—as well as 10 unidentified compounds in this same chemical class. The specific blend of these and other carboxylic acids varied between different high attractive subjects. Therefore, there may be more than one way for a person to be highly attractive to mosquitoes. We did not identify any compounds that were reproducibly enriched on the skin of the least attractive humans, consistent with the idea that these individuals lack mosquito attractants, rather than emitting a shared set of repellent compounds. A limitation of our study is that the chemical derivatization we used specifically focused on metabolites containing carboxylic acid groups. However, a strength of this approach is that this aqueous-compatible derivatization allowed us to measure metabolites with a wide range of volatilities using a single analytical method, from highly volatile short chain fatty acids to medium volatility acids to longer chain fatty acids of lower volatility. A previous study that analyzed interindividual differences in mosquito attractiveness focused on different classes of compounds and identified five—6-methyl-5-hepten-2-one, octanal, nonanal, decanal, and geranylacetone—that were enriched on the skin of weakly attractive humans (*39*). Individually these compounds reduced mosquito flight activity and/or attraction, suggesting that some people may release natural repellents that make them less attractive to mosquitoes. One of our subjects, Subject 90, had high levels of carboxylic acids in their skin emanations but was only weakly attractive to mosquitoes. It is plausible that Subject 90 produces higher levels of a natural repellent that would counteract the elevated levels of carboxylic acids, but this was not tested in our study.

### Mosquito attractiveness of a given person is stable over years

Our study focused on humans who were empirically determined to lie at extremes of the mosquito-attractiveness spectrum. Several subjects were tested regularly over the entire 3-year duration of the study, allowing us to determine that extreme mosquito attraction is remarkably stable. All humans tested in our study were at least somewhat attractive to mosquitoes when their skin odor blend was presented as a single stimulus. However, we estimate that skin odor from Subject 33 was over 30 times as potent that of the least attractive human sampled, Subject 28. Our behavioral assay results corroborate anecdotal evidence that context matters for how attractive a person is to mosquitoes in a real-world setting, since mosquitoes feed opportunistically. If one human walks into a highly mosquito-infested environment alone, they may receive many bites, regardless of their overall attractiveness level because they are the only feeding option. Mosquito preferences matter more in group settings. The “mosquito-magnet” in the group may receive the most bites, leaving the less attractive humans largely untouched. We observed that mosquitoes show preferences between individuals who are extremely highly attractive on their own. For example, nylons from Subjects 31, 24, and 33 were all similarly highly attractive when competed against an unworn nylon. However, Subject 33 was more attractive than the two other subjects when competed head-to head in the two-choice assay. Furthermore, mosquito preferences were transitive: Subject 33>Subject 25>Subject 28. This suggests that mosquitoes distinguish the scent of two human samples using cues that exist along a continuum. Consistent with this idea, we were able to dilute the dose of nylon from a highly attractive human to the point where it was equally attractive as a nylon from a low attractor human. We propose that exceptionally high or low attractiveness to mosquitoes is a “fixed” trait, caused by factors that remain constant over a period of several years, even when environmental factors are not strictly controlled. We did not require study participants to maintain a constant diet, exercise regimen, and we did not limit changes to medication or personal care product usage over the 3 years of the study. It has been shown that identical twins are more similarly attractive to mosquitoes than fraternal twins (*18*), suggesting a genetic component to mosquito attractiveness. Moreover, the blend of carboxylic acids that characterizes individual human body odor types is more similar in monozygotic twins than unrelated subjects (*70*). We speculate that genetically determined skin characteristics and/or other very stable inter-individual differences contribute to making someone highly or weakly attractive to mosquitoes.

### Sensory mechanisms of mosquito discrimination between humans

In this study, we discovered that mutating any one of the three carboxylic acid-sensing IR co-receptors, *Ir8a*, *Ir76b*, or *Ir25a*, strongly reduced overall attraction to human scent, but did not abolish the ability of any of these mutants to discriminate between highly and weakly attractive people. We also found that elevated levels of skin-derived carboxylic acids – ligands that activate the IR pathway – are associated with high mosquito attractiveness. Although our data do not allow us to conclude that skin carboxylic acid abundance causes an individual to be highly attractive to mosquitoes, the association between these phenotypes is interesting. In contrast, *Orco* mutants, which have an impaired ability to sense other classes of chemical compounds, including esters, ketones, alcohols, and aldehydes, are highly attracted to humans (*5, 60, 61*). In our behavioral assays using *Orco* mutants, there was no significant difference in either activation or attraction, or in their ability to distinguish between any of the pairs of human subjects.

Because no single mutation was able to disrupt the ability of mosquitoes to discriminate between two subjects, we speculate that there is extensive redundancy in the detection of human-derived skin odors. This redundancy may be due to central olfactory coding mechanisms or to the recently described co-expression of ORs and IRs in the same olfactory sensory neuron (*63, 71*). One possible mechanism of IR redundancy could lie with their use of three and not one co-receptor as for the OR system. Removing any one IR co-receptor might have only a partial effect on the ability of the mosquito to detect odorants sensed by IRs. One possibility is that the three IR co-receptors together with ligand-selective IRs collaborate to tile the chemical space of carboxylic acids, such that removing any single IR co-receptor reduces the overall attraction to humans but allows the mutants to retain the ability to detect differences in levels of carboxylic acids.

Our findings argue against the idea that mosquitoes distinguish between highly and weakly attractive humans using a single odor, such as lactic acid, as has been suggested (*13*). If this were the case, we would expect that *Ir8a* mutants, which cannot sense lactic acid (*61*), would lose their preference for highly attractive humans. Instead, we propose that mosquito magnets produce elevated levels of multiple mosquito-attractant compounds, and that this drives mosquito preferences. Among the chemical entities we found to be elevated on highly mosquito attractive human skin, we identified 3 compounds as straight chain, unsaturated carboxylic acids with 15,17, and 19 carbons. The relative contribution of these 3 identified compounds and the unidentified entities to mosquito preferences remains unknown. Notably, fatty acids with greater than 10 carbons are not very volatile, so it is unclear whether the compounds we identified are directly sensed by the mosquito, or whether they may give rise to more volatile components that are also enriched on the skin of mosquito magnet subjects.

### Microbiota influence over human skin acid production

Free fatty acids are more abundant on the skin surface of humans than non-human animals (*72*), and therefore they may be an important indicator to mosquitoes that there is a human nearby. Human skin is unique among mammals because it has relatively little hair and numerous eccrine sweat glands across most of its surface. Some animals, including humans, produce a specialized waxy substance from sebaceous glands called sebum. In humans, sebum is triglyceride-rich, thereby producing a characteristic surface lipid composition, which contains about 25% free fatty acids (*72*). The unique skin lipid composition of humans is thought to have protective effects, such as limiting sun damage in the absence of protective hair, and emulsifying eccrine sweat, preventing its overly rapid evaporation to allow for appropriate body temperature regulation (*73, 74*). Human skin acids are astonishingly diverse, with branched, odd-chain, and esterified fatty acids reported, along with skin-specific patterns of desaturation (*73*). This complexity is also likely reflected in the 10 unidentified features upregulated in highly attractive subjects, as many appeared to be structural isomers of the identified straight-chain acids. Given the vast array of acid types found on the skin, it is unlikely that two individual humans will possess the same exact complement of acids in the exact same ratios, potentially giving each human a unique chemical signature. Skin bacteria contribute to the pool of free fatty acids found on human skin by producing several types of fatty acid synthetase enzymes that allow them to produce diverse types of acids themselves (*75, 76*) and by cleaving free fatty acids from human sebum triglycerides using lipase enzymes (*72*). Additionally, recent work has shown that skin microbiota composition is remarkably stable within an individual over time, even though skin is exposed to a constantly fluctuating environment (*77*). Most viable skin bacteria reside in the pores, where they are protected from external factors, such as hygiene habits and seasonal weather changes (*78*). It is reasonable to think that an individual’s skin microbiota contributes to their skin acid composition, which we have shown to be remarkably stable over time.

### Closing remarks

The attraction preferences of disease-vectoring mosquitoes have important public health implications, since it is estimated that in disease endemic areas a small fraction of humans is more frequently targeted, and these individuals serve as a reservoir of pathogens (*11, 79*). Understanding the mechanistic basis for mosquito biting preferences will suggest new ways to reduce mosquito attraction to humans and curb the spread of dangerous arboviruses. Studies in humans (*29–31, 33*) and mice (*32*) have demonstrated that infection with malaria parasites alters their attractiveness to mosquitoes by altering the odors they produce, leading to greater pathogen transmission. Understanding what makes someone a “mosquito magnet” will suggest ways to rationally design interventions such as skin microbiota manipulation to make people less attractive to mosquitoes. We propose that the ability to predict which individuals in a community are high attractors would allow for more effective deployment of resources to combat the spread of mosquito-borne pathogens.

## ACKNOWLEDGMENTS

We thank Lindy McBride, Jessica Zung, and members of the Vosshall Lab for comments on the manuscript; all study participants for their generous participation in this research; Gloria Gordon and Libby Mejia for expert mosquito rearing; Ben Matthews and Veronica Jové for discussion and guidance on *Ir76b* and *Ir25a* mutagenesis strategies; Maya Davidov and Naquia Unwala for help with pilot behavioral studies to establish the feasibility of this work; Arka Rao, Sara Nunes Violante, Michelle Saio, Ruben Jose Jesus Faustino Ramos, and members of the Donald B. and Catherine C. Marron Cancer Metabolism Center for guidance with metabolomic experiments and GC/QTOF-MS data analysis; and the PIT Crew: Jim Petrillo, Peer Strogies, and Dan Gross of the Rockefeller University Precision Instrumentation Technologies resource center for advice on assay fabrication.

## AUTHOR CONTRIBUTIONS

M.E.D. carried out or supervised all experiments and data analysis in the paper with additional contributions from co-authors. T.M. generated and carried out initial characterization of *Ir76b* and *Ir25a* mutants. L.C.D. helped conceive the preference study in Figure 1E and carried out all behavioral experiments in the paper except the validation cohort experiments in Figure 7B, which were carried out by E.V.Z. M.E.D. conducted the initial GC/QTOF-MS experiments in Figure 6. D.J.B. together with M.E.D. carried out the validation GC/QTOF-MS experiments in Figure 7. C.S.J. contributed statistical expertise and code for analysis of both mosquito behavior data and untargeted analysis of GC/QTOF-MS data. J.R.C. supervised D.J.B. and provided technical and conceptual guidance for the GC/QTOF-MS experiments. M.E.D. and L.B.V. together conceived the study, designed the figures, and wrote the paper with input from all authors.

## Materials and Methods

### Human and animal ethics statement

Blood-feeding procedures and mosquito behavior with live human hosts were approved and monitored by The Rockefeller University Institutional Review Board (IRB protocol LVO-0652) and the Rockefeller University Institutional Animal Care and Use Committee (IACUC protocol 20068-H). Human subjects gave their written informed consent to participate in this study.

### Human subject information

64 healthy human subjects participated in this study. Age at inception of study on 12/7/2017: mean 29.8, median 29, range 19-57 years. Self-identified gender: 37 female, 26 male, 1 non-binary. Due to sample size limitations, we intentionally did not subdivide these groups further to investigate the contribution of demographic factors such as sex, age, and ethnicity, or behavioral factors such as diet, personal care, or activity levels on mosquito attractiveness, which we determined empirically. The behavior experiments in Figures 1-5, and Figure 8 included only subjects from the initial cohort (Subjects 19, 24, 25, 28, 30, 31, 32, 33). Seven of these subjects were tested in GC/QTOF-MS Experiments 1.1-1.4 in Figure 6. Subject 24 had moved away before we performed the experiments in Figures 5-7. These 7 subjects from the initial cohort (Subjects 19, 25, 28, 30, 31, 32, 33) were also analyzed about a year later alongside 56 newly recruited subjects (Figure 7B). The full single stimulus olfactometer behavior dataset for all 63 subjects is available on Zenodo (DOI: 10.5281/zenodo.5822539), including data for the 18 subjects whose data are plotted in Figure 7B, comprising 4 subjects from the initial cohort (Subjects 19, 28, 31, and 33) and 14 subjects from the validation cohort. These 18 subjects were selected for inclusion in GC/QTOF-MS Experiments 2.1-2.4, based on their overall level of attractiveness to mosquitoes in initial screening experiments (a subset of the data presented in (Figure 7B), and their availability to participate in additional behavior and GC/QTOF-MS experiments.

### Human skin odor collection

Human subjects washed their forearms with Dove unscented soap and water, dried them with clean laboratory paper towels, and then wore nylons sleeves on both forearms between the wrist and elbow for 6 hours. Nylon sleeves were prepared by using scissors to remove 2” of fabric from the tip of the stocking foot of knee highs (L’eggs brand Everyday, Amazon), so that the modified stocking was open on both ends. Subjects wore 2 nylon sleeves on each arm: a brown experimental nylon was worn next to the skin, and a black outer nylon was placed over the experimental nylon to minimize contamination of the inner nylon. Subjects wore nylons during the day and were allowed to perform typical daytime activities but were asked not to exercise or drink alcohol while wearing the nylons. After the 6-hour wearing period, nylons were deposited in Whirl-Pak bags and kept at -20°C for 1-10 days before behavioral and chemical analysis. For the “round-robin” two-choice assay experiment in Figure 1E, we competed 2 nylon sleeves that had been worn on the same day by 2 different human subjects, to strictly control the “age” of the nylons being used to determine if mosquitoes preferred one subject over the other. We relaxed the requirement that subjects wear the nylons on the same day for subsequent two-choice assay experiments, since small differences in nylon age did not change the preference of mosquitoes for specific human subjects. In Figures 2,3,5, and 8, we compared nylons that had been worn within 2 days of each other by two different subjects. This allowed us to better accommodate the schedules of human subjects, who were not always available on the same day over the extended duration of the study.

### Mosquito rearing and maintenance

*Aedes aegypti* wild-type laboratory strains (Orlando and Liverpool) were reared in an environmental room maintained at 70-80% relative humidity and 25-28°C, as previously described (*60*). All animals were maintained with a photoperiod of 14 hours light: 10 hours dark throughout larval, pupal, and adult life stages. Adult mosquitoes were provided constant access to 10% sucrose. Female mosquitoes were fasted for 14-24 hours without sucrose in the presence of water prior to behavioral experiments. For stock maintenance, females were blood fed on live mice.

### *Ir25a* and *Ir76b* mutant strain generation

*Ir25a* and *Ir76b* mutants were generated using methods described previously (*80*). sgRNA sequences were designed with the CRISPOR v4.3 sgRNA design tool (http://crispor.tefor.net/) (Concordet, 2018) using the following parameters: Genome, *Aedes aegypti* – yellow fever mosquito – NCBI GCF_002204515.2 (AaegL5.0); Protospacer Adjacent Motif (PAM), 20bp-NGG -Sp Cas9, SpCas9-HF1, eSpCas9 1.1. For each gene, two pairs of sgRNA with predicted MIT Specificity Scores ≥95 were selected for targeted double stranded break-induced mutagenesis, with each pair flanking roughly 250 base pairs within exon 2 for *Ir76b* and exons 2 and 3 for *Ir25a*. sgRNA DNA templates were prepared by annealing oligonucleotides as previously described using the following target sequences:

Ir25a-sgRNA1: GTTGAGCTACTAACCGTCGA

Ir25a-sgRNA2: TACTGACAGCAAAGGGCTGT

Ir25a-sgRNA3: CCTACGGTTTCCGCATCAAC

Ir25a-sgRNA4: AAGAAGGCGACTTGAGGCAA

Ir76b-sgRNA1: GTTACACCGAACGTCAGAA

Ir76b-sgRNA2: TACTCTGGTCGGACGCGGTG

Ir76b-sgRNA3: CTCCTTTCAATCGGGACGTG

Ir76b-sgRNA4: CAACGGCCAGCAGCGATACC

*In vitro* transcription was performed using HiScribe Quick T7 kit (NEB E2050S) following the manufacturer’s protocol. Following *in vitro* transcription and DNAse treatment for 15 minutes at 37°C, sgRNA was purified using RNAse-free SPRI beads (Ampure RNAclean, Beckman-Coulter A63987), and eluted in Ultrapure water (Invitrogen, 10977–015). To further facilitate isolation of loss-of-function mutants, 200 bp single-stranded DNA oligodeoxynucleotide (ssODN) donors were designed as a template for homology-directed repair (*80*). The ssODN donor had homology arms of 88–90 bases on either side of the inner most sgRNA target sites (sgRNA2 and sgRNA3 for both *Ir25a* and *Ir76b*), flanking an insert with stop codons in all three frames of translation and a *BamHI* restriction site (IDT). Since homology arms contained sgRNA target sites for the outer most sgRNA target sites (sgRNA1 and sgRNA4 for both *Ir25a* and *Ir76b*), PAM motifs for the respective sgRNAs were mutated to avoid ssODN donor cleavage. The sequences for each ssODN follow below with left and right homology arms italicized, stop codons underlined, and *BamHI* restriction site highlighted in bold:

Ir25a-ssODN:

TTGCGCTGAACTATATAAGAAAGAACCCAAGCCTCGGACTTTCAGTTGAGCTACTAACCG TCGAATGAAACCGTACTGACAGCAAAGGGC<u>TAATAA</u>**GGATCC**A<u>TAA</u>C<u>TAA</u>GGAACCGGCC AAGAAGGCGACTTGAGGCAATAGCGATCTCTATCAAACGTAAAAAGCAACTATCTGTTGCAGGTTTGCTACAAAACTTTA

Ir76b-ssODN:

TCTAATTGCATCGAACTCTCTTTCCCACGTTCAACAGGACTGGCCGCTGAGTTACACCGA ACGTCAGAATAGTACTCTGGTCGGACGCG<u>TAATAA</u>**GGATCC**A<u>TAA</u>C<u>TAA</u>GGGTGTGGATT TTGATTCTGGTATCGCTGCTGGCCGTTGGTCCAATCATCTACGGAATGCTGATTGTGCGGTACAAAATGACCAAAGACAA

For each target gene, approximately 500 wild-type *Aedes aegypti* (Liverpool LVP-IB12 strain) embryos were injected with a mixture containing recombinant Cas9 protein (PNA Bio, CP01) at 300 ng/µl, 4 sgRNAs at 40 ng/µl each, and donor ssODN at 125 ng/µl. Embryos were injected by the Insect Transformation Facility at the University of Maryland Institute for Bioscience & Biotechnology Research. Embryos were hatched and G0 females were crossed to wild-type Liverpool males, and their G1 offspring were screened for germline mutation by PCR amplification and Sanger sequencing the regions flanking the sgRNA target sites. Two unique stable mutant lines, each resulting in an early stop codon due to a frameshift mutation, were isolated from each injection. For *Ir25a*, one isolated mutant line had a 209-bp deletion with an ssODN integration (*Ir25a^BamHI^*), and the other had a 19-bp deletion (*Ir25a^19^*). For *Ir76b*, one isolated mutant line had a 61-bp deletion (*Ir76b^61^*), and the other had a 32-bp deletion (*Ir76b^32^*). Virgin females from each mutant line were backcrossed to wild-type Liverpool males for 8 generations prior to establishment of stable homozygous lines. To control for potential CRISPR-Cas9 off-target effects on behavior, homozygous mutant lines were intercrossed to generate heteroallelic mutants that were tested in all behavior experiments alongside appropriate genetic controls. It was previously shown that although *Ir76b* mutant *Anopheles coluzzii* females show normal attraction to human host cues, they fail to blood feed and therefore produce no offspring (*81*). We also observed deficits in blood-feeding and egg-laying in homozygous *Aedes aegypti Ir25a* and *Ir76b* strains, albeit far less severe than the *Anopheles coluzzii Ir76b* mutants. All homozygous *Ir25a* and *Ir76b* mutant strains were maintained by blood-feeding on a human arm. We determined empirically that these mutant mosquitoes fed more avidly when they were 14 days post eclosion, rather than 7 days, which is the standard timepoint at which we blood feed to propagate other strains. These strains did not feed effectively on live mice or on an artificial membrane blood feeder. A volunteer inserted an arm into a standard BugDorm rearing cage (30 cm^3^) cage for 30 minutes at ambient temp/humidity. These cages contained a mixed population of males and females, which had been allowed to mate freely since eclosion. Mosquitoes had access to the entire hand and forearm of the subject. For context, normal strains feed to repletion on a human arm after 5-10 minutes, even when the arm is placed outside of the cage netting and not inside the cage. Feeding of these mutants was unsuccessful when the human arm was placed against the netting on the outside of the cage. About 50% of females were engorged at the end of a given 30-minute feeding session, while the rest were partially fed or unfed. Notably, a given female took much longer to probe the arm repeatedly before successfully feeding. No attempt was made to refeed females that did not feed during a given 30-minute feeding session. Four days after feeding, an oviposition cup was placed into the cage and as many eggs as possible were collected over a 4-day period. After this stage the entire process was repeated once or twice with the same cage to obtain enough eggs to propagate the strains and carry out experiments. It is our impression that *Ir25a* females but not *Ir76b* females laid fewer eggs than wild type, but we did not investigate this further in the course of this study. Unlike the observed difficulties with the homozygous mutants, heterozygous mutants fed normally and laid normal numbers of eggs.

### *Ir25a* and *Ir76b* mutant genotyping

Genotypes were confirmed using Phire Tissue Direct PCR Master Mix (Thermo Fisher, F170L) followed by gel electrophoresis and Sanger sequencing (Genewiz) using the following primer combination for each mutant allele:

*Ir25a^BamHI^*, *Ir25a^19^*

Forward: AATACTTGAGGAGTCGTTGAAT

Reverse: GAAGCAATGCCTTGTACTTATG *Ir76b^61^*

Forward: AGCCGAATATGAAGGTCAAGC

Reverse: CAGCACCTGTTCCTTGTCTT *Ir76b^32^*

Forward: TGCATCGAACTCTCTTTCCC

Reverse: CGATAGCTAAGATGCCAGTACAT

The *Ir25a^BamHI^* allele was detected by either a 159 bp deletion or presence of an exogenous *BamHI* restriction site from the donor ssODN. *BamHI* restriction digest of the PCR product generated a single 764 bp fragment in wild-type animals and two fragments (275 bp and 329 bp) in the *Ir25a^BamHI^* mutant. For mutants with small deletions, the presence or absence of endogenous restriction enzyme target sites was used to distinguish between mutant and wild-type alleles. PCR products were generated and digested with the indicated enzyme, producing the indicated bands in mutant and wild type:

*Ir25a^19^* with *MspI*

Wild type: 502 and 262 bp; *Ir25a^19^*: 764 bp

*Ir76b^61^* with *BstUI*

Wild type: 367 and 397 bp; *Ir76b^61^*: 764 bp

*Ir76b^32^* with *BstNI*

Wild type: 392 and 365bp; *Ir76b^61^*: 757 bp

All genotyping experiments were performed with a no DNA control as well as fragment size validation using 1Kb Plus DNA ladder (ThermoFisher Scientific, 10787026). See Supplementary Figure S1.

Given the severe host-seeking and blood-feeding deficits displayed by *Ir25a* and *Ir76b* female homozygous mutants, it is difficult to maintain these as homozygous strains. For the benefit of scientists wishing to work with these new strains, we have devised a crossing scheme that uses heterozygous mutant females to propagate the mutant alleles. An important aspect of this approach is that *Ir25a* and *Ir76b* mutants do not carry a fluorescent marker at their respective gene loci. This strategy consists of first crossing homozygous *Ir25a* or *Ir76b* mutant males to heterozygous females from the corresponding *Ir25a-QF2* and *Ir76b-QF2* gene-sparing knock-in driver lines (*71*). These strains contain the *3xP3-dsRed* marker integrated at the *Ir25a* or *Ir76b* genetic locus. This initial cross will generate both fluorescent and non-fluorescent heterozygous mutants at a 1:1 ratio. Then, taking the fluorescent heterozygous mutant females, these animals will again be crossed to the homozygous mutant males. This cross will generate non-fluorescent homozygous mutants and fluorescent heterozygous mutants at a 1:1 ratio. At this point, mutant alleles can easily be maintained by collecting eggs from the cross between non-fluorescent homozygous mutant males with fluorescent heterozygous females isolated at each generation. Since the mutations and the locations of the inserted fluorescent markers are tightly linked, we expect the recombination rates between the mutation sites and the markers to be extremely rare. However, occasional genotyping is recommended to ensure proper propagation of each of the mutant alleles.

### Behavioral assays

The single choice olfactometer assay referred to in this work (Figure 4C, Figure 7B) is the same assay previously referred to as the “Quattroport” in an earlier publication (*65*), because the assay allows 4 independent single stimulus olfactometer trials to be run in parallel. In this work, we refer to the Quattroport assay as the “single stimulus olfactometer assay” to avoid confusion about the number of stimuli being presented to each group of mosquitoes in a single trial. We repurposed components of the Quattroport assay to create the two-choice olfactometer assay, which allows us to compete 2 different stimuli against each other in the same trial. We performed two separate preference trials in parallel, increasing throughput over the Gouck olfactometer assay (*64*). Details of fabrication and operation of the two-choice assay are available on Zenodo (DOI: 10.5281/zenodo.5822539). Air flow and CO_2_ conditions for the two-choice olfactometer assay were carried as described for the Quattroport (*65*). Human forearm-worn nylon sleeves were used as the stimulus in all behavior figures with the two-choice olfactometer assay, except Figure 1B in which human subjects placed their forearm over a hole in the stimulus box lid, exposing 12.9 cm^2^ of skin to mosquitoes (demonstrated with a mannequin arm in Figure 1A). We shuffled the order in which different stimuli were assessed over the course of the day, and we randomized the position of different stimuli across all stimulus boxes of the assays, to reduce time of day and position effects. All behavioral experiments were carried out in an environmental room set to 25°C, 70-80% relative humidity. The day before behavior was measured, 20 female mosquitoes (aged 7-14 days post-eclosion, mated) were sorted under cold anesthesia into each start canister and given access only to water for 18-22 hours. The same canisters were used for the single stimulus and two-choice assays. However, twice as many females were tested in each two-choice assay trial (40 females per trial, 20 in each of 2 canisters) as in each single stimulus trial (20 females per trial). In both assays, mosquitoes were acclimated to a carbon-filtered air stream for 10 minutes (Donaldson Ultrac-A). CO_2_ was then introduced into the air stream for 30 seconds, at which point mosquitoes were released, and given 5 minutes to assess the stimulus or stimuli. Mosquitoes were prevented from contacting stimuli by a mesh divider. Sliding doors between assay compartments allow the experimenter to count the number of mosquitoes that were not activated, or activated but not attracted, or both activated and attracted to the stimulus. In the single stimulus assay, mosquitoes that entered an attraction trap were scored as attracted to the stimulus. In the two-choice assay, mosquitoes that entered a cylindrical flying tube or the adjacent trap, both downwind of the stimulus, were scored as attracted to that stimulus. In both assays, all mosquitoes that left the start canister were scored as activated. We are confident that mosquitoes use olfactory information to detect nylon stimuli, since mosquitoes cannot contact or taste the nylon in our assays due to the presence of a mesh barrier, and there are no visual cues that differentiate nylons worn by different subjects.

### Cleaning behavioral assay apparatus

Between trials, the assay apparatus was vacuumed to remove live and dead mosquitoes, and air was flowed through the assay for 5-10 minutes to flush out residual CO_2_ and odor. Two-choice assay parts were cleaned before use in every experiment, as described below. In some cases, assay parts needed to be cleaned between trials, so that they could be re-used during the same behavior experiment. Whenever possible, we constructed enough replicate assay parts to reduce the need to wash parts between trials, which slows down assay throughput. Experimenters always wore gloves when cleaning and handling clean assay parts. Detailed procedures are described here for how and when we cleaned each part of the two-choice assay (going from left to right in the schematic in Figure 1A: 1) start canisters, 3D-printed connector joints, and accompanying acrylic sliding doors were washed in a dishwasher (Miele Optimal Series dishwasher, Cascade Original “Actionpacs” detergent pods) at least 2 days before behavior and allowed to air dry completely before female mosquitoes were loaded into the canisters on the day before behavior; 2) the flying box was too big to be washed in the sink or dishwasher, so the inside of the box was sprayed down with 70% ethanol from a laboratory spray bottle and this was wiped down with laboratory paper towels, and allowed to air dry; 3) two cylindrical flying tubes were washed in the dishwasher as described above and allowed to air dry; 4) Acrylic stands that support the cylindrical flying tubes were washed in the dishwasher as described above; 5) the complete supply of attraction traps and accompanying 3D printed joins and acrylic sliding doors were washed in the dishwasher as described above at least one day before the experiment and allowed to air dry. When we performed more trials than we had traps available, we hand washed the traps used in the first few trials using hot water and soap (Bac Down Handsoap, Decon Labs, Inc., Catalog #: 7001), and allowed them to air dry before re-using them in trials later in the day. 6) For stimulus boxes and lids, the cleaning procedure was the same as that for the traps. The complete supply of stimulus boxes and lids was washed in the dishwasher as described above at least one day before the experiment and allowed to air dry. When we performed more trials than we had stimulus boxes available, we hand washed the stimulus boxes used in the first few trials using hot water and soap and allowed them to air dry before re-using them in trials later in the day. 7) The 3D printed stop piece which connects the air/CO_2_ supply to the stimulus box was wiped down with 70% ethanol that had been sprayed onto a laboratory paper towel before the first trial, and between every trial. Similarly, the single stimulus olfactometer assay parts were cleaned before every experiment. This means that, going from left to right in the schematic in Figure 4C: 1) start canisters, 3D-printed connector joints, and accompanying acrylic sliding doors were washed in a dishwasher at least 2 days before behavior and allowed to air dry completely before female mosquitoes were loaded into the canisters on the day before behavior; 2) the two cylindrical flying tubes were hand washed in the sink with hot water and soap and a bottle brush (Dr. Brown’s, Amazon, ASIN: B01NCUKCC0), and allowed to air dry ; 3) Acrylic stands that support the cylindrical flying tubes were washed in the dishwasher as described above; 4) the complete supply of attraction traps and accompanying 3D printed joins and acrylic sliding doors were washed in the dishwasher as described above at least one day before the experiment and allowed to air dry. When we performed more trials than we had traps available, we hand washed the traps used in the first few trials using hot water and soap and allowed them to air dry before re-using them in trials later in the day. 5) For stimulus boxes and lids, the cleaning procedure was the same as that for the traps. The complete supply of stimulus boxes and lids were washed in the dishwasher as described above at least one day before the experiment and allowed to air dry. When we performed more trials than we had stimulus boxes available, we hand washed the stimulus boxes used in the first few trials using hot water and soap and allowed them to air dry before re-using them in trials later in the day. 6) The 3D printed stop piece which connects the air/CO_2_ supply to the stimulus box was wiped down with 70% ethanol that had been sprayed onto a laboratory paper towel before the first trial, and between every trial.

### Behavior inclusion criteria

Two-choice olfactometer assay data indicating the overall percent mosquito activation and attraction in response to a single human subject stimulus, are presented in Figure 2A, Figure 3A, and Figure 5A for all control trials performed (i.e. all trials examining Subject 25 vs Subject 25, before the application of any inclusion criteria). In Figure 2B, Figure 3B, and Figure 5B, we report the percent of control trials that met the inclusion criteria used in later parts of these Figures, which required that there were: 1) at least 30 live mosquitoes at the end of the assay and 2) at least 10 mosquitoes attracted to either stimulus. We chose these inclusion criteria because IR mutants displayed large defects in overall attraction to human subjects, and we wished to examine preferences specifically using trials which passed a minimal threshold of overall attraction to human odor. Trials with very low overall attraction to either stimulus, could give misleading results, because they are subject to “jackpotting” effects. For example, if 2 mosquitoes were attracted to Subject A, and 1 mosquito was attracted to Subject B, this is not a meaningful difference in preference, so we would exclude this trial for having <10 animals attracted to either subject. To assay mosquito preferences between genotypes in Figure 2C-D, Figure 3C-D, and Figure 5C-E, we present only trials which passed the inclusion criteria. We discarded trials in which substantially fewer mosquitoes were loaded into the assay (unintentionally), and which very few mosquitoes (<10) were attracted to either stimulus. There were some exceptions to the application of inclusion criteria, described here: the experiment in Figure 1C compared mosquito preferences between several stimulus pairs that were expected to (and did) result in very low levels of attraction. These stimuli were: 1) unworn nylons versus no stimulus, and 2) no stimulus versus no stimulus. For this experiment, we did not require that more than 10 mosquitoes were attracted to either stimulus. We only required that there were >30 live animals at the end of the trial. For single stimulus olfactometer experiments (Figure 4D-G, Figure 7B), we included trials with >14 live mosquitoes at the end of the trial.

### Nylon sleeve behavioral assay stimuli

For human odor host-seeking assays, a 7.62 cm x 10.16 cm piece of the brown experimental nylon was cut and used as a stimulus in either the two-choice olfactometer assay or the single stimulus olfactometer assay. This was the largest size of material that could be laid flat on the bottom of the stimulus boxes used in both assays. By using this size of nylon stimulus, we could cut 3 pieces of nylon from each nylon sleeve, yielding a total of 6 pieces of nylons from each day of nylon wearing, or enough for 6 trials of that subject. Experimenters always handled nylon sleeves with gloves and cleaned scissors with 70% ethanol between samples. Subjects wore 2 nylon sleeves on each of their 2 forearms (one brown experimental nylon next to their skin, covered by one black protective nylon, as described above). We asked subjects to remove both nylon sleeves they wore on each arm as a single “unit” (keeping the black protective nylon outside the brown experimental nylon) and to place these in a Whirl-Pak bag (Nasco, Catalog #: B01062) in the freezer. This allowed us to determine which side of the brown experimental nylon had touched the subject’s skin (the side that was not touching the black outer nylon), and we marked this side with a fine point permanent marker. The side of the nylon that touched the subject’s skin was placed facing upwards in the stimulus box, to ensure consistency between trials. In all behavior experiments, nylon samples were de-identified before being cut and presented to mosquitoes in the behavioral assay (either the two-choice assay or the single stimulus olfactometer assay), such that experimenters were blinded to the Subject ID. Each piece of nylon was attached to a flexible plastic rectangle of the same size (cut with scissors from plastic file folders: Letter size, Office Depot, Catalog #: 700259), using small binder clips (3/4” size, Office Depot Catalog #: 808857). The color of the plastic rectangle indicated the de-identified label given to that nylon, corresponding to the source of the nylon. For instance, in a given experiment, Subject 33 was designated as Subject “A” by someone other than the experimenter, and then the experimenter cut the de-identified nylon pieces and attached all Subject “A” nylons to red plastic cards. The color of the card was chosen randomly for each experiment, and colors rotated between subjects. Each nylon piece was used for only one trial and then discarded. Binder clips and plastic cards were washed with soap and water as described above and set out to air dry at the end of each experiment.

### Statistical analysis of behavior

Statistical analyses were performed using R. Two-choice assay data were analyzed by comparing the percent of mosquitoes attracted to each of two stimuli, for several pairs of stimuli, using Wilcoxon rank-sum tests with Bonferroni correction (Figure 1B-E, Figure 2C-D, Figure 3C-D, Figure 5C-E). Nonparametric effect size (ES) was calculated as ES= Z/sqrt(N) where N is the number of observations (Figure 1E) (*82*). A power analysis was used to pre-determine the approximate sample size needed for Figure 1E (G*Power software). Wilcoxon rank-sum tests with Bonferroni correction were used to compare the percent of mosquitoes in each of three categories (attracted, activated but not attracted, or not activated) between wild type and each mutant genotype in Figure 2A, Figure 3A, Figure 5A. Single stimulus olfactometer assay data in Figure 4D-G were analyzed by using a Kruskal-Wallis test to compare the percentage of mosquitoes activated or attracted across 4 genotypes: including wild-type controls, 2 heterozygous mutants, and the heteroallelic null mutants. When indicated, post hoc analysis used pairwise Wilcoxon rank-sum tests with Bonferroni correction to compare wild-type mosquitoes to each of the other 3 genotypes for a given stimulus (Figure 4D-G).

### Calculation of the “Attraction Score”

The attraction score reported in Figure 1F was calculated as follows. For each of the 28 subject pairs in the “round-robin tournament”, we calculated the total number of mosquitoes attracted to each subject in the pair, by adding the number that were attracted across every trial performed for that specific pair in Figure 1E. For example, for the Subject 33 vs Subject 28 pair, we summed the total number of mosquitoes attracted to Subject 33 or to Subject 28 across the 18 trials in which they were compared to each other: Subject 33 attracted 468 mosquitoes, whereas Subject 28 attracted only 35 mosquitoes. We took the difference between these values (468-35=433) and divided it by 18 trials to get the average “margin of victory” per trial for this subject pair (433/18=24). Since Subject 33 attracted 24 more mosquitoes than Subject 28 per trial, on average, Subject 33 was “awarded” 24 points, and Subject 28 was awarded 0 points. Subject 33 also attracted more mosquitoes than the remaining 6 other subjects, with average “margin of victory scores” of 15, 18, 20, 21, 23, and 23. The “attraction score” for each subject, reported in Figure 1F, represents the sum of 7 average “margin of victory” values. In the example above, the “attraction score” is calculated by summing the average “margin of victory scores” for Subject 33, compared to the 7 other subjects: 15+18+20+21+23+23+24=144.

### Sample preparation for gas chromatography/quadrupole time of flight-mass spectrometry (GC/QTOF-MS)

Nylon sleeves that had been stored at -20°C for 1-10 days were thawed and cut into 5.08 cm x 2.54 cm pieces. Experimenters always handled nylon sleeves with gloves and cleaned scissors with 70% ethanol between samples. For each subject, 6 separate pieces (5.08 cm x 2.54 cm each) of the same nylon sleeve were analyzed in each of the 4 experiments shown in Figure 6 (Experiments 1.1-1.4), and 5 pieces of the same nylon sleeve were analyzed in each of the 4 experiments shown in Figure 7 (Experiments 2.1-2.4). Extraction solvent was 80% methanol (CID: 67-56-1, Methanol, Fisher Chemical, Catalog #:A456-4) prepared with Millipore water and spiked with deuterated isotope-labeled internal standards sourced from Cambridge Isotope Laboratories: nonanoic acid-D17 (CID: 130348-94-6, Catalog #: DLM-9501-0.5), propanoic acid-D5 (CID: 60153-92-6, Catalog #: DLM-1919-5), phenol-D5 (CID:4165-62-2, Catalog #:DLM-695-1), acetate-D4 (CID:1186-52-3, Catalog #: DLM-12-10 ), butyrate-D7 (CID: 73607-83-7, Catalog #: DLM-1508-5) and valerate-D9 (CID: 115871-50-6, Catalog #:DLM-572-1). Each piece of nylon was extracted in 1 mL of extraction solvent in a 5 mL Eppendorf tube (Fisher Scientific Catalog #: 14-282-301). 5 mL Eppendorf tubes containing extraction solvent and nylon were kept overnight at -20°C. Then, each tube was vortexed for 20 seconds to ensure thorough extraction of the nylon. A P1000 pipet with standard 1000 µL pipet tip was used to press the nylon against the side of the 5 mL Eppendorf tube, so that the extraction solvent was squeezed out of the nylon. Before doing this, the plunger of the pipet had been pressed down, so that it could be released after squeezing the nylon, to pick up the extraction solvent, before it could be reabsorbed into the nylon sleeve. Typically, 700-900 µL of nylon “extract” (a cloudy mixture of extraction solvent plus nylon compounds) was recovered from each sample, and this was immediately transferred to a 2 mL glass GC vial (Wheaton µL MicroLiter autosampler vials; 12×32mm, Catalog #: 11-1200) and secured with a septum cap.

### Pentafluorobenzyl bromide (PFB-Br) derivatization

100 µL of the liquid nylon extract was manually transferred using a pipet to a new glass autosampler vial, containing 100 µL of borate buffer (100mM boric acid CID: 10043-35-3, Sigma Aldrich, Catalog #: 339067, adjusted to pH 10 using 10 M NaOH (CID: 1310-73-2, Fisher Chemical, Catalog #: S318500. Subsequent steps were performed using a single needle liquid handling system (Gerstel MPS). First 400 µL of 100 mM pentafluorobenzyl bromide (PFB-Br, CID: 1765-40-8, Sigma-Aldrich, Catalog number: 101052), prepared in acetonitrile (Fisher Chemical; Catalog #: A955-4) and 400 µL cyclohexane (Sigma-Aldrich, Catalog #: 650455-4L) were added sequentially to the reaction vials. Samples were then sealed using crimp caps (Wheaton MicroLiter Catalog #: 11-0040A) and heated to 65°C with shaking for 1 hour to esterify carboxyl and hydroxyl groups (Supplementary Figure S3A). Samples were allowed to cool to room temperature and centrifuged for 2 minutes at 2,000 rpm, 20°C to promote phase separation, then returned to the robotic sample preparation instrument. Finally, the upper cyclohexane layer was transferred to clean autosampler vials (Wheaton MicroLiter, Catalog #: 11-1200) and 50 µL further transferred into a second clean autosampler vial pre-filled with 450 µL cyclohexane, creating a 10-fold dilution of the derivatized nylon extracts that was subsequently used for GC/QTOF-MS analysis.

### GC/QTOF-MS data collection

GC/QTOF-MS data was collected using an Agilent 7890B gas chromatograph (GC) and an Agilent 7200 quadrupole time-of-flight (QTOF) mass spectrometer (MS), fitted with a Gerstel MPS Robotic Autosampler. 1 µL of the sample volume was injected in splitless mode onto a DB5ms column (30 m × 250 μm, 0.25 μm film thickness; Agilent Technologies, Catalog #:19091S-433). GC conditions were as follows: initial oven temperature 60°C (hold 1 minute), ramp at 25°C/minute to 300°C (hold 2.5 minutes), ramp at 120°C/minute to 60°C (hold 1 minute); the total run time was 16.1 minutes. The inlet, transfer line, chemical ionization (CI) source, and quadrupole temperatures were set to 280°C, 300°C, 150°C, and 150°C respectively. The emission current was set to 4.2 µA and data collection was in 2 GHz mode, mass range *m/z* 50-650. For all GC/QTOF-MS experiments, the sample injection order was randomized using the “RAND” function in Microsoft Excel. The same GC column was used for data acquisition shown in Figure 6 and Figure 7, column trimming as part of routine maintenance was responsible for slightly shorter retention times shown in Figure 7. The Q-TOF mass spectrometer was operated in negative chemical ionization (nCI) mode with methane as the reagent gas (1 mL/minute); in nCI mode, derivatized molecules undergo electron capture dissociation (ECD) and are detected as deprotonated (M-H) ions.

### GC/QTOF-MS data analysis

The workflow for GC/QTOF-MS data analysis is summarized in Supplementary Table S1. Raw data (Agilent “.D” files) were analyzed using Agilent MassHunter software. Untargeted feature finding was performed using Agilent Unknowns Analysis software (v10.1) (Supplementary Table S1, step 1). Specifically, data was first converted using SureMass peak detection (absolute height filter >1000 counts, RT window size factor=50, Extraction window +-50ppm, use base peak shape=yes, sharpness threshold 25%, Ion peaks minimum=1, maximum=1). Subsequently, feature finding was performed on all worn nylon samples across 4 replicate experiments for Experiments 1.1-1.4 (Figure 6) and, again for Experiments 2.1-2.4 (Figure 7). Putative features, denoted as accurate mass @ retention time, were exported from Unknowns Analysis as a .csv file, and imported into R. A custom deduplication script was used to remove most “fuzzy” duplicate features with similar mass and retention time values, and features detected in <10 data files, resulting in a list of 1494 features for the experiments in Figure 6, and 1925 features for the experiments in Figure 7 (Supplementary Table S1, step 2). This feature list was imported into Agilent Quantitative Analysis software (v10.1) and a targeted data re-extraction performed (settings: +Gaussian smoothing, +area filter ≥ 0, RT window: ± 0.05 minutes extraction window ± 100ppm) (Supplementary Table S1, step 3), resulting in a fully aligned dataset across 4 replicate experiments. Raw peak areas were then exported as a .csv file, and a custom R script was again used to remove features present in <10% of samples, impute missing values with 1/2 the lowest value for that feature, and log2 transform data (Supplementary Table S1, step 4). The resulting .csv file was imported into Agilent Mass Profiler Professional software (v15.1). We considered high quality features to be those found in at least 50% of samples in at least 1 subject group, with a coefficient of variation <40% in at least 1 subject group. Putative human skin-derived features were considered those at least 2-fold enriched (false discovery rate, FDR<0.05) in any Subject’s worn nylons versus unworn nylons and solvent blank controls (Supplementary Table S1, step 5). Volcano plots were used to filter on features differentially abundant in the high versus low attractor subjects in each individual experiment of 4 replicate experiments (Supplementary Table S1, step 6; Figure 6D, Figure 7D). Venn diagrams were used to identify hits that met the differential abundance criteria in high versus low attractors across all 4 replicate experiments (Supplementary Table S1, step 7): Experiments 1.1-1.4 (Figure 6E) and Experiments 2.1-2.4 (Figure 7E). Redundant features with closely matching mass and retention times, that were missed by the earlier deduplication step were removed manually, after inspecting the raw data in Agilent Qualitative Analysis software (Supplementary Table S1, step 8). Data File S1 contains lists of mass and retention times for unknown features that were found in 4 replicate experiments, either Experiments 1.1-1.4 or Experiments 2.1-2.4. For all targeted data re-extractions, 100 ppm mass accuracy was used to extract the detected pseudomolecular [M-H]^-^ion, unless the peak was considered saturated, in which case the [(M+1)-H]^-^ isotope was used consistently across all data files. For Experiments 1.1-1.4, we calculated the median abundance of each compound in each of 4 experiments for each of 2 high attractor subjects and 2 low attractor subjects (Supplementary Figure S2B). Then we compared these values for each compound between the high and low attractor groups using a nonparametric linear mixed effects model for repeated measures based on ranks, with group as a fixed effect and subject as a random effect. Benjamini-Hochberg correction was applied for multiple comparisons (FDR<0.1). For Experiments 2.1-2.4, we calculated the median abundance value for each compound in each subject across 4 replicate experiments, and then we compared these values between the high (n=11) vs low attractor (n=7) groups (Figure 7H) using a Wilcoxon rank-sum test. Benjamini-Hochberg correction was applied for multiple comparisons (FDR<0.1). Statistical analysis was performed in R version 4.0.5. Heatmaps were generated using Metaboanalyst (v5.0; https://www.metaboanalyst.ca/) (*83*) using autoscaled data, clustering was Ward’s linkage and Euclidean distance measure.

### GC/QTOF-MS compound identification

To identify compounds of interest, we used the accurate mass of the molecular ion (M-H) to predict chemical formulas using Agilent Qualitative Analysis software (v10.0). Formula prediction was constrained to C, H, O, N atoms, and formulas within 100 ppm mass error of the measured mass were considered (Supplementary Table S1, step 9). Predicted formulas for unknown compounds are included in Data File S1. To achieve positive identification, we spiked nylon samples with authentic standards (details below) and asked whether the area of the unknown peak increased symmetrically (Supplementary Table S1, step 10). Overlaid extracted ion chromatograms generated in Agilent Qualitative Analysis software for 3 hit compounds are shown in Supplementary Figure S3G. Commercially standards used were: propanoic acid (CID: 5818-15-5, Millipore Sigma; Catalog #: 94425-1ML-F), butyric acid(CID: 107-92-6; Millipore Sigma; Catalog #: 19215-5ML), pentanoic acid (CID: 109-52-4; Millipore Sigma; Catalog #: 75054-1ML), hexanoic acid(CID: 142-62-1; Millipore Sigma; 21529-5ML), heptanoic acid (CID: 111-14-8; Millipore Sigma; Catalog #: 43858), octanoic acid (CID: 124-07-2; Millipore Sigma; Catalog #:21639), nonanoic acid (CID: 112-05-0; Millipore Sigma; Catalog #:73982), decanoic acid (CID: 334-48-5; Millipore Sigma; Catalog #:21409), undecanoic acid (112-37-8; Millipore Sigma; Catalog #:89764), dodecanoic acid (143-07-7; Millipore Sigma; Catalog #:61609), tridecanoic acid (CID: 638-53-9; Millipore Sigma; Catalog #: 91988), tetradecanoic acid (CID: 62217-71-4; Millipore Sigma; Catalog #:70079), pentadecanoic acid (CID: 1002-84-2; Millipore Sigma; Catalog #:91446), hexadecanoic acid (CID: 57-10-3; Millipore Sigma; Catalog #:76119), heptadecanoic acid (CID: 506-12-7; Millipore Sigma; Catalog #:H3500), octadecanoic acid (CID: 38003-60-0; Millipore Sigma; Catalog #:85679), nonadecanoic acid (CID: 646-30-0; Millipore Sigma; Catalog #:72332), icosanoic acid (CID: 506-30-9); Millipore Sigma; Catalog #: 39383).

## DATA AVAILABILITY

Supplementary Figures S1-S3 and Supplementary Table S1 accompany the paper. All raw data in the paper and instructions for the behavioral assays are available on Zenodo (DOI: 10.5281/zenodo.5822539) at this link: https://zenodo.org/record/5822539#.YdXST2jMIuU.

---

**Supplementary Figure S1 - Related to Figure 4.**
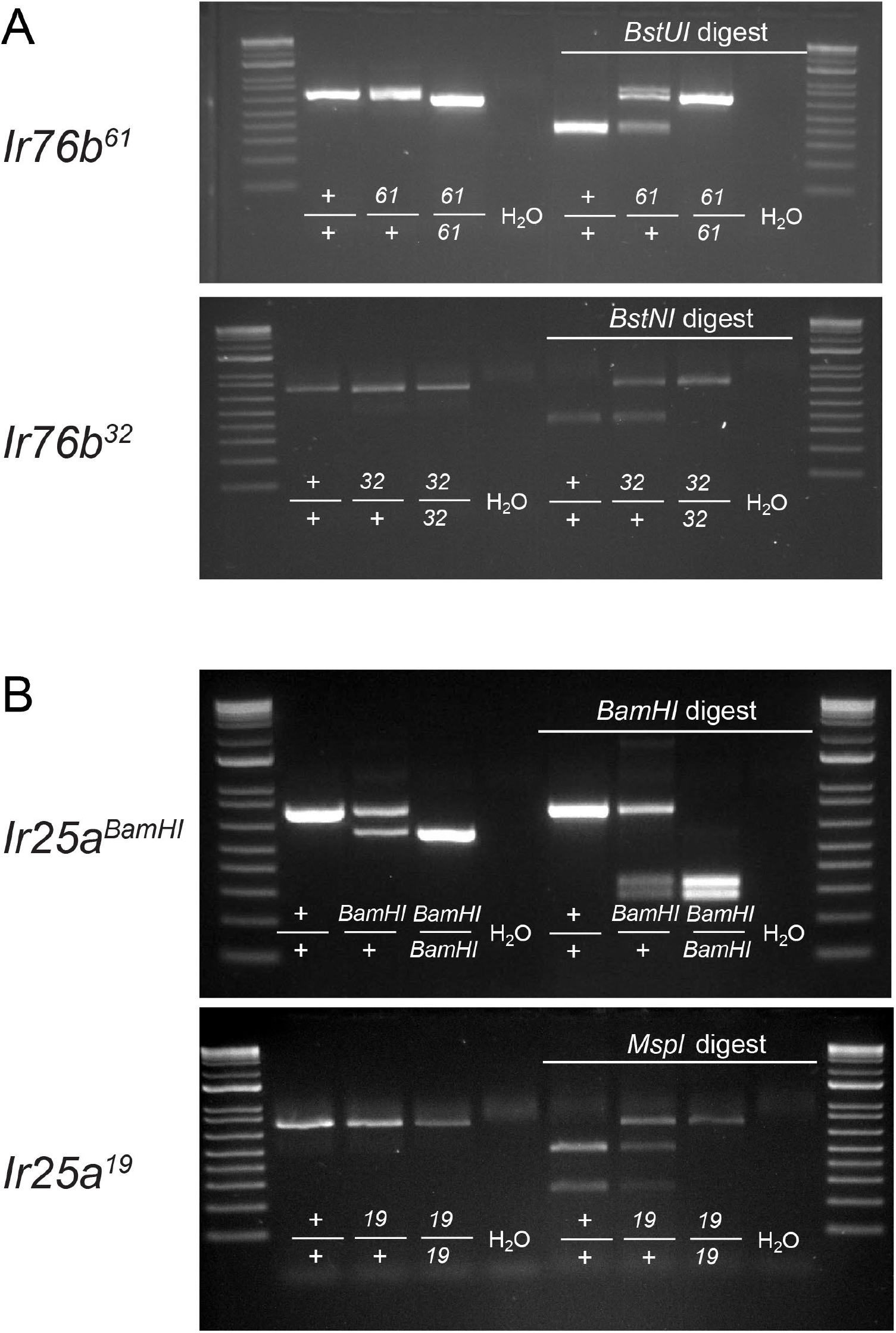
PCR genotyping *Ir76b* and *Ir25a* mutant strains. (**A-B**) Agarose gel electrophoresis images of PCR fragments before (left) and after (right) restriction with the indicated restriction enzyme to genotype the indicated *Ir76b* (A) and *Ir25a* (B) mutants.

**Supplementary Figure S2 - Related to Figure 6 and Figure 7.**
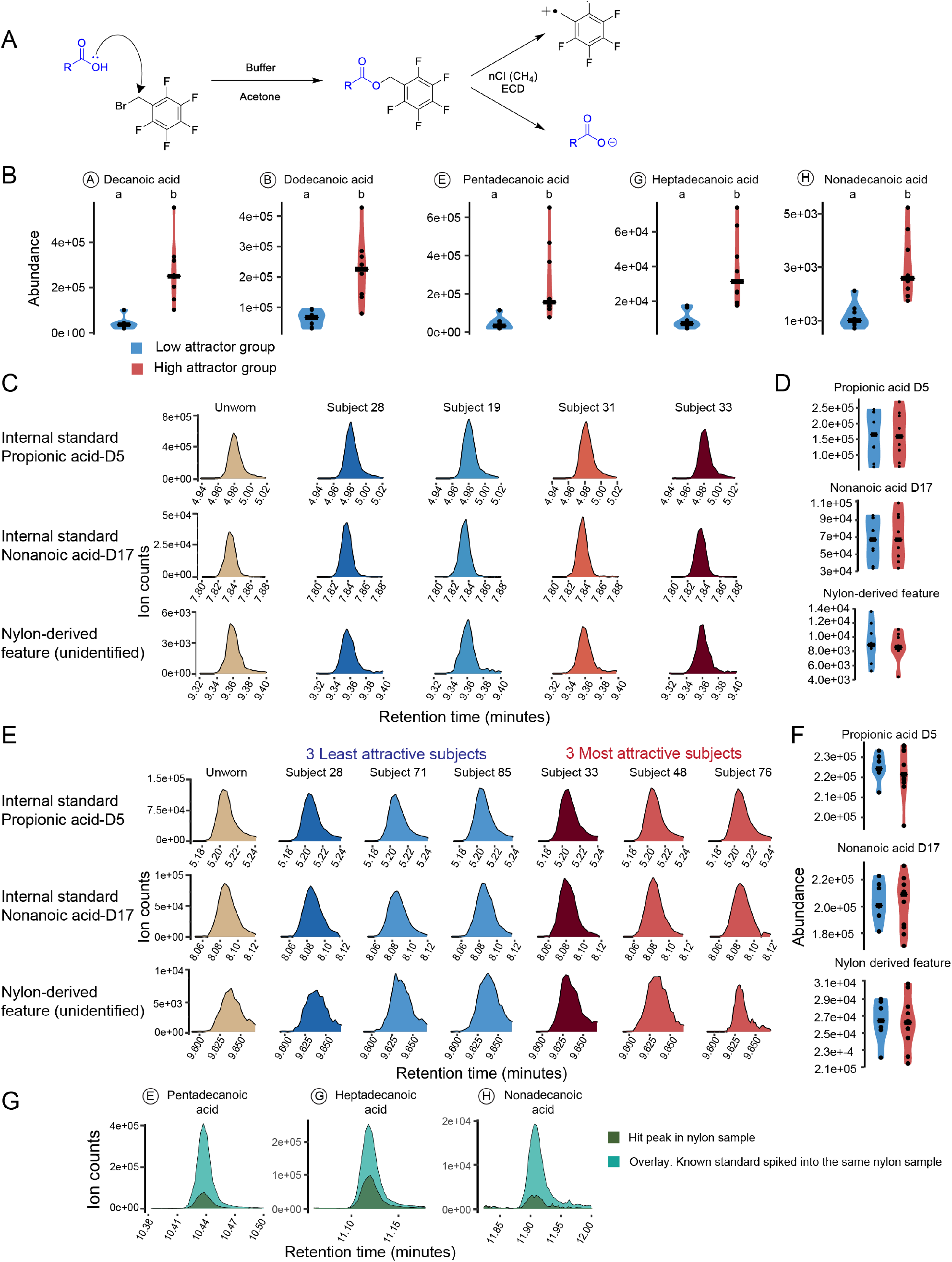
Additional description and validation of GC/QTOF-MS data. **(A)** Mechanism of pentafluorobenzyl-bromide (PFB-Br) derivatization reaction. **(B)** Abundance of several carboxylic acids in low attractors (Subjects 19, 28) vs. high attractors (Subjects 31,33). Each dot represents abundance of the indicated compound for one subject in one experiment (median of 4 replicate samples). Nonparametric linear mixed-effects model followed by Benjamini Hochberg FDR correction (p<0.1) was used. Violins labeled by different lowercase letters were significantly different. In addition to the 5 compounds plotted here, 3 additional compounds were also found to significantly more abundant in the high attractors than low attractors: tetradecanoic acid, hexadecenoic acid, and icosanoic acid. Tridecanoic acid was not significantly different. **(C)** Representative extracted ion chromatograms (EICs) for 3 control compounds: 2 deuterated internal standards and 1 nylon-derived compound, from the indicated human subjects in Experiment 1.1. **(D)** Abundance of control compounds shown in (C). Each dot represents the abundance (median of 4 replicate samples) of the indicated compound for one subject in 1 of 4 experiments: Experiments 1.1-1.4). **(E)** Representative extracted ion chromatograms (EICs) for 3 control compounds: 2 deuterated internal standards and 1 nylon-derived compound, from the indicated human subjects in Experiment 2.1. **(F)** Abundance of control compounds shown in E. Each dot represents the abundance (median of 4 replicate samples) of the indicated compound for one subject in 1 of 4 experiments: Experiments 2.1-2.4). **(G)** Extracted ion chromatograms of three identified hit compounds, overlaid with extracted ion chromatograms of the same sample spiked with a known standard.

**Supplementary Figure S3- Related to Figure 6 and Figure 7.**
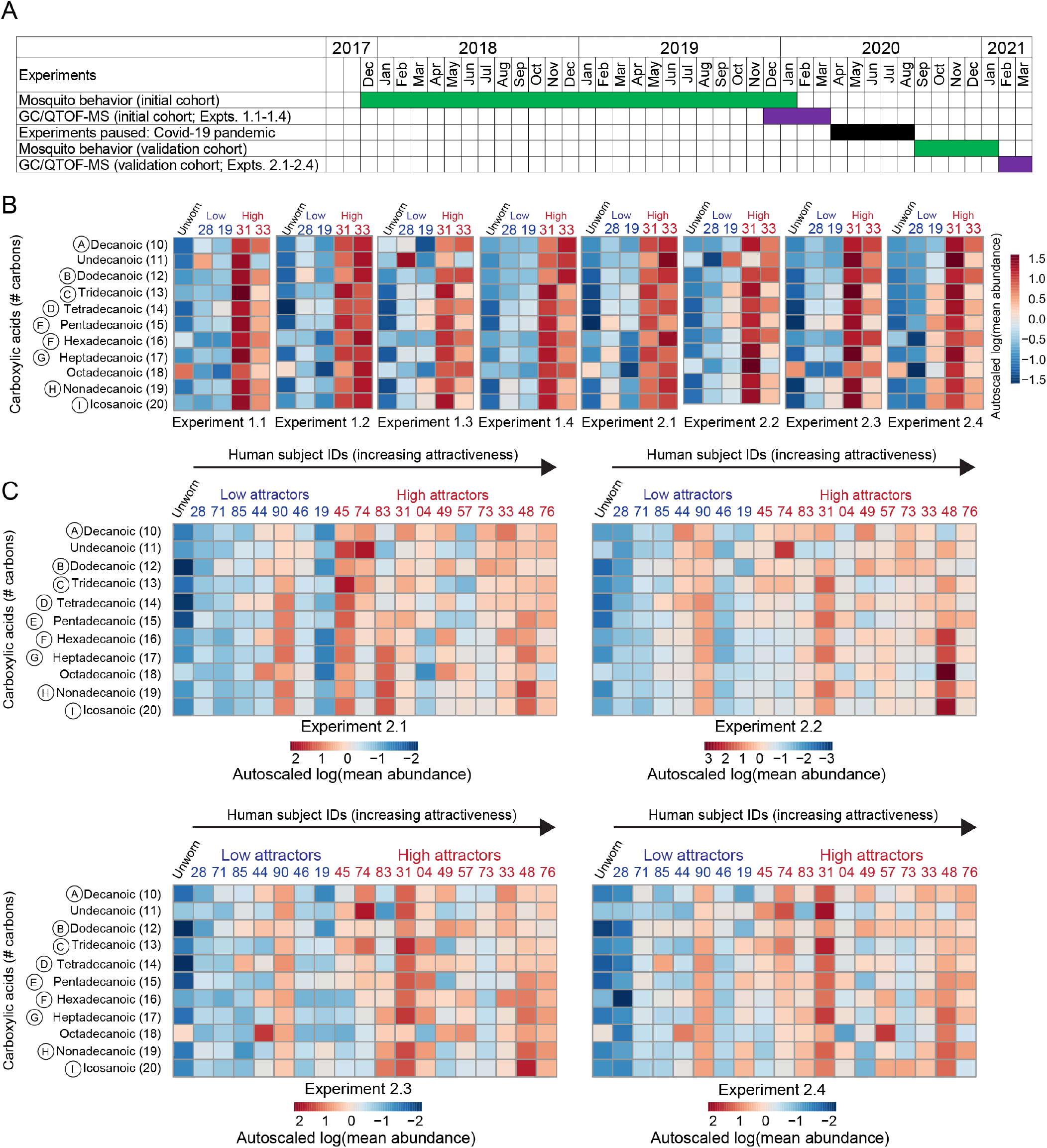
Intra-individual stability of skin chemistry. **A)** Timeline of behavior (green blocks) and GC/QTOF-MS experiments (purple blocks), relative to one another. 64 human volunteers participated in this study, and nylons from 18 of these were analyzed by GC/QTOF-MS. Nylons from four of these subjects (Subjects 19, 28, 31, 33) were repeatedly tested behaviorally over a 3-year period and also analyzed using GC/QTOF- MS in 2 sets of 4 replicate experiments (Experiments 1.1-1.4, Experiments 2.1-2.4) that were conducted 1 year apart. (B) Heatmaps depicting quantified abundance of carboxylic acids with 10-20 carbons, averaged across 4 replicate samples per experiment, in 4 subjects. Each heatmap represents one of 8 independent experiments. Experiments 1.1-1.4 were conducted about a year before Experiments 2.1-2.4. (C) Heatmaps depicting quantified abundance of carboxylic acids with 10-20 carbons, averaged across 5 replicate samples per experiment, in 18 subjects from the validation cohort. Each heatmap represents one of 4 independent experiments, conducted about a week apart.

**Supplementary Table 1:**

| Supplementary Table S1 |  |  |
| --- | --- | --- |
| Untargeted workflow for GC/QTOF-MS |  | # features at each step |
| Description of analysis step | Experiments 1.1-1.4 | Experiments 2.1-2.4 |
| 1. Feature finding performed on all worn nylon samples (from 4 experiments) – using Agilent Unknowns Analysis software – Sure Mass deconvolution | ~133,000 | ~300,000 |
| 2. Initial feature deduplication (R code “step 2”) | 1,494 | 1,925 |
| 3. Targeted analysis of deduplicated feature list in all samples (Agilent Qualitative Analysis 10) | 1,494 | 1,925 |
| 4. Using R code “step 4”: replaced zero values with NA, remove features present in <10% of all samples, impute missing values with 1/2 the lowest value for that feature, perform Log2 transformation | Not determined | Not determined |
| 5. Imported data back into Agilent Mass Profiler Professional software: baselining: “none”, scaling: “none”, and then use a fold change filter to select high quality features in each experiment<br>a. found in ≥50% of samples in ≥1 subject group<br>b. coefficient of variation ≤40% in ≥1 subject group<br>c. 2-fold upregulated (FDR< 0.05) in ≥1 subject group vs. unworn and vs. solvent control group | Exp. 1.1: 404<br>Exp. 1.2: 376<br>Exp. 1.3: 345<br>Exp. 1.4: 378<br>(Note: 204 features were found in 4 of 4 experiments) | Exp. 2.1: 633<br>Exp. 2.2: 619<br>Exp. 2.3: 604<br>Exp. 2.4: 597<br>(Note: 161 features were found in 4 of 4 experiments) |
| 6. Filter on “hits”: features differentially abundant in high attractor vs low attractor subject: ≥1.5-fold change (FDR<0.1) (Agilent Mass Profiler Professional – Volcano plot)<br>Note: In Exp. 1.1-1.4: 2 high attractor subjects: 31 & 33 were compared 2 low attractor Subjects 19 & 28. (Subject 24 was not available for GC/MS analysis)<br>Note: In Exp. 2.1-2.4: 7 low attractor subjects were compared to 11 high attractor subjects | Exp. 1.1: 210<br>Exp. 1.2: 219<br>Exp. 1.3: 147<br>Exp. 1.4: 240<br>(Figure 6D shows Experiment 1.1) | Exp. 2.1: 130<br>Exp. 2.2: 106<br>Exp. 2.3: 100<br>Exp. 2.4: 105<br>(Figure 7D shows Experiment 2.3) |
| 7. Filter on hits found in 4 of 4 experiments (Agilent Mass Profiler Professional – Venn diagram) | 93 | 23 |
| 8. Manually remove redundant features from CSV file in Microsoft Excel | 51 (Figure 6E table, more info: Zenodo*) | 13 (Figure 7E table, more info: Zenodo*) |
| 9. Formula prediction (Agilent Qualitative Analysis 10) | 42 (more info: Zenodo*) | 5 (more info: Zenodo*) |
| 10. Positive identification of hit compounds (matched to known standard) | 9 | 3 |

\*Zenodo (DOI: 10.5281/zenodo.5822539) at this link: https://zenodo.org/record/5822539#.YdXST2jMIuU

